# Genome admixture among four hare species in Iberia: focus on the broom hare (*Lepus castroviejoi*)

**DOI:** 10.1101/2024.07.23.604737

**Authors:** João Souto, João P. Marques, Liliana Farelo, José Costa, João Queirós, Christian Pietri, Fernando Ballesteros, Paulo C. Alves, Pierre Boursot, José Melo-Ferreira

## Abstract

Pleistocene climatic fluctuations have often driven range shifts and hybridization among related species, leaving present-day genomic footprints. In the Iberian Peninsula, *Lepus timidus*, after its post-deglaciation retreat, has left extensive mitochondrial DNA traces in three other hare species, but the genomic correlates and underlying biogeographic scenarios are still incompletely understood. This study focuses on *Lepus castroviejoi*, endemic to the Cantabrian region, using its non-Iberian sister species, *L. corsicanus*, for comparison. By analyzing coalescent patterns from 10 genomes, we estimate that these species remained isolated since their divergence, around 50,000 years ago, consistent with their current allopatry. Further analyses with 25 additional genomes indicate that small fractions of the *L. castroviejoi* genome originate from *L. granatensis*, *L. timidus*, and *L. europaeus* (0.72%, 0.08%, and 0.04%, respectively). Introgression dating based on tract lengths suggests *L. granatensis* was already admixed with *L. timidus* when it hybridized with *L. castroviejoi*, which could explain the *granatensis*-*timidus* ancestry tract junctions detected in *L. castroviejoi*. Genomic segments with such junctions contain genes enriched for cell signaling and olfactory receptor activity, possibly facilitating genetic exchange. This research demonstrates how genomic ancestry inferences can reveal complex multiway admixture histories and illuminate past biogeographic events.

## Introduction

Modern analyses of complete genomes from natural populations have highlighted that the gene pools of species are frequently shaped by genetic exchanges with other species, at different stages of their divergence process (Moran et al., 2021). The continuous impact of introgressive hybridization over the long course of the evolution of species leaves layered genetic signatures that persist throughout their diversification process (Ferreira et al., 2021). While the determination of which parts of the genome do not participate in such exchanges may inform about the speciation process, identifying the regions that are extensively exchanged among species can inform about the potential role of introgression in adaptation (Edelman & Mallet, 2021; Aguillon et al., 2022), as exemplified in butterflies (Pardo-Diaz et al., 2012), hares (Jones et al., 2018; Giska et al., 2019) or sunflowers (Whitney et al., 2010). However, pervasive introgression can also arise from demographic processes. In scenarios of range replacement accompanied by hybridization, introgression can become prevalent at the invasion front due to allele surfing (Currat et al., 2008; Seixas et al., 2018). Introgression patterns therefore reflect a combination of biogeographic events and selective pressures that shaped the evolution of species, involving transient or persistent interactions, range changes or range replacements (Rougeux et al., 2019; Villa-Machío et al., 2023). As a consequence, the gene pool of extant species is often a mosaic of different ancestries (Edelman & Mallet, 2021), which mark the history of range changes and interspecific contacts. Characterizing historical introgression events can therefore be an important mean of reconstructing the biogeographic history of species (e.g. Seixas et al., 2018).

Hares (*Lepus* spp.) from western Europe have been extensively affected by introgressive hybridization during their evolutionary history. Their present distribution ranges are the result of recurrent range changes, overlaps and hybridization events since late Pleistocene (Alves et al., 2008b; Melo-Ferreira et al., 2012; Seixas et al., 2018). The mountain hare (*Lepus timidus*) currently ranges from northern Europe to Far East Russia, and is present in some isolated populations, such as in Ireland, Great Britain and the Alps (Angerbjörn, 2018), but palaeontological records show that it was present much further south at the last glacial maximum, reaching the Iberian Peninsula (Altuna, 1970; Lado et al., 2018). Its subsequent northward retreat resulted from the post-glacial climate warming and possibly also from the invasion of temperate species (Melo-Ferreira et al., 2007; Acevedo et al., 2015). These range changes caused multiple contacts with other species, accompanied by hybridization, since mitochondrial DNA (mtDNA) introgression from *L. timidus* is now found in all species presently thriving in the Iberian Peninsula: the Iberian hare (*Lepus granatensis*), distributed all over most of Iberia; the broom hare (*Lepus castroviejoi*), endemic to the Cantabrian Mountains; and the European brown hare (*Lepus europaeus*), present in the northern fringe of the Peninsula, and extending towards central Europe (Melo-Ferreira et al., 2011; Seixas et al., 2018). In the case of *L. granatensis*, geographical variation of nuclear introgression across the Peninsula indicates that hybridization occurred during south-north expansion of this species, replacing the resident *L. timidus* after the last glacial maximum (Marques et al., 2017; Seixas et al., 2018). Ancient DNA analysis revealed that this northward expansion reached southern France, followed by a range contraction back to Iberia, likely driven by the arrival of *L. europaeus* (Lado et al., 2018), establishing a hybrid zone in the north of the Peninsula (Melo-Ferreira et al., 2014). In Italy, the Corsican hare (*L. corsicanus*) is present in the Apennines, and has recently diverged from its sister species *L. castroviejoi*, present in the Iberian Peninsula (Alves et al., 2008a; Ferreira et al., 2021). Discordances between mitochondrial DNA and consensus nuclear DNA phylogenies have suggested that in these two species the mitochondrial genome was replaced by that of *L. timidus*, as a result of introgressive hybridization events, presumably involving their common ancestor (Melo-Ferreira et al., 2012). Here, we analyse whole genome sequencing data from *L. castroviejoi* and *L. corsicanus*, model their divergence history, and use inferences of introgression to reconstruct their biogeographic dynamics in the context of the known history of the other western European hares.

## Methods

### Sampling, sequencing and variant calling

We used whole-genome sequences generated for 5 *L. castroviejoi*, 5 *L. corsicanus*, 10 *L. granatensis*, 10 *L. europaeus*, 4 *L. timidus* and 1 *L. americanus*. These datasets contained new and previously published data (Seixas, 2017; Seixas et al., 2018; Giska et al., 2019) (Figure 1; see Table S1 for a detailed account of sample and data origin). The samples are part of CIBIO-InBIO’s collection and were kindly provided by researchers or donated by hunters during permitted hunting season. No animal was killed for this research. All samples were collected before the Nagoya protocol came into force where applicable.

**Figure 1.**
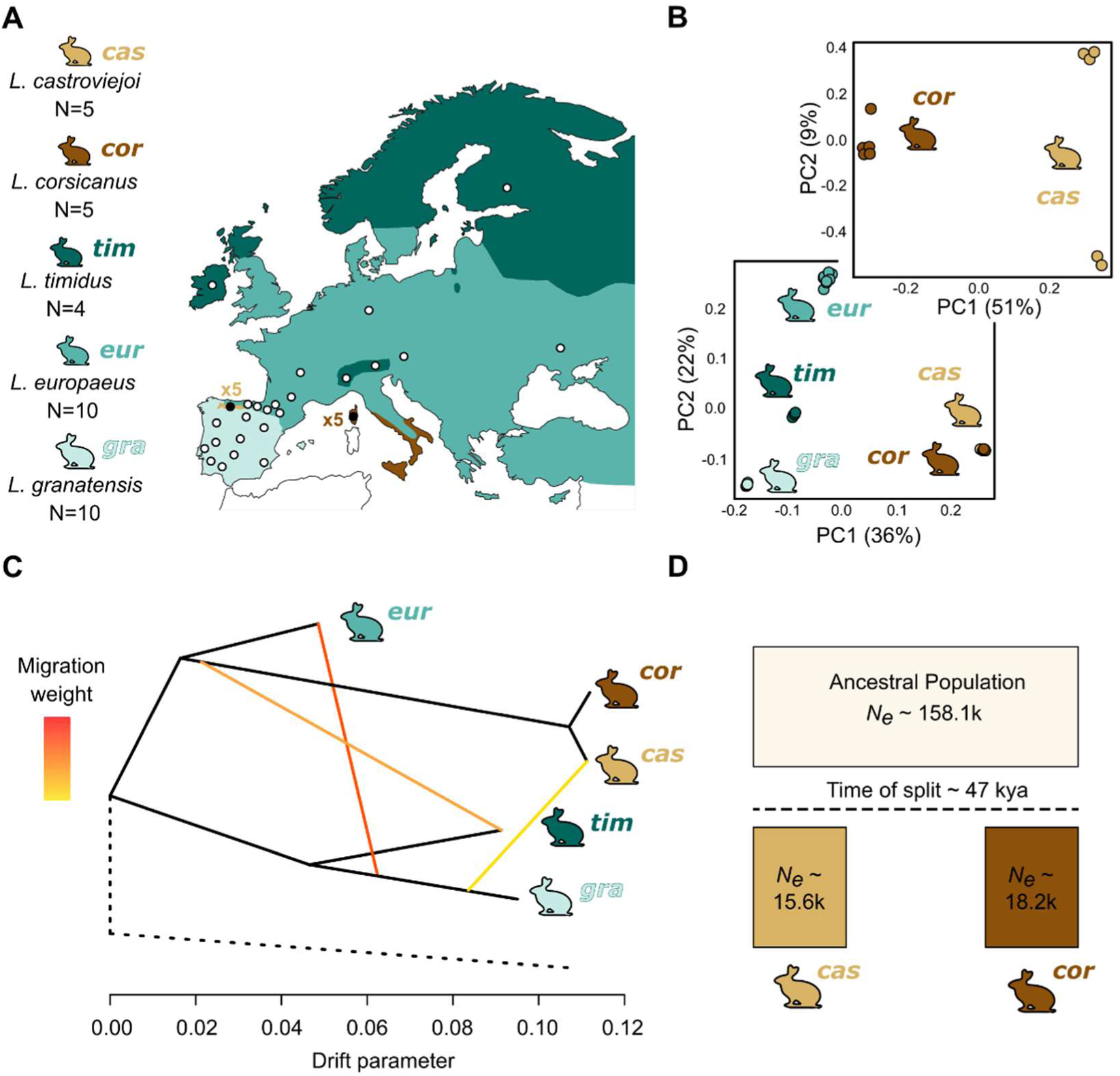
Sampling, species structure and demographic inferences. **A.** Distribution in Western Europe of the hare species analysed in this work (based on Mitchell-Jones et al., 1999), and sampling localities represented by circles. **B.** Principal Component Analysis (PCA) based on SNP data of European hare species (2,999,121 SNPs; bottom panel) and for *L. castroviejoi* and *L. corsicanus* (433,825 SNPs; top panel). **C.** Species relationships inferred from allele frequency data using Treemix (dashed line indicates the outgroup branch, *L. americanus*) with 3 ingroup migration events. **D.** Demographic history of divergence between *L. castroviejoi* and *L. corsicanus* inferred with G-PhoCS, based on a model without post-split gene flow (*N_e_*, effective population size; kya, thousand years ago; k, thousand individuals).

Genomic DNA was extracted using the JETquick Tissue DNA Spin Kit (GENOMED) from ear or internal organ tissue, which were originally preserved in ethanol or RNAlater. Illumina TruSeq DNA PCR-free genomic libraries were prepared and sequenced using an Illumina HiSeq 1500 sequencer at the NEWGEN sequencing platform at CIBIO (Vairão, Portugal), generating paired-end sequence data (2x100-125 bp).

The first 5 bases and adapters from the end of raw reads were removed with Cutadapt version 1.8 (Martin, 2010), and low quality bases were trimmed using Trimmomatic v0.33 (Bolger et al., 2014) (quality < 20 at the end of reads, and 4 consecutive bp with average quality < 30). A hare pseudo-reference genome was built based on our whole genome sequencing dataset using an iterative mapping approach (Sarver et al., 2017) and using the European rabbit (*Oryctolagus cuniculus*) reference genome as template (OryCun2.0). Filtered reads were mapped to this pseudo-reference using the BWA-MEM algorithm with default parameters (Li & Durbin, 2009). Read pairs were corrected and sorted using Samtools v1.3 (Li et al., 2009). Insertion-deletions (indels) were realigned using the Genome Analysis Toolkit (GATK v3.2-2) (McKenna et al., 2010; DePristo et al., 2011) and duplicated reads were removed with Picard Markduplicates (http://broadinstitute.github.io/picard/). Finally, Bcftools 1.10.2 mpileup (Li, 2011) was used to perform the multi-sample SNP/genotype calling for each species independently, with minimum base and mapping qualities of 20. The resulting VCF files were merged, indels were removed, and repetitive regions annotated using the European rabbit genome (from https://genome-euro.ucsc.edu/) were excluded with Bcftools 1.10.2.

### Genetic structure and relationships among hare species from Europe

To assess how the genetic variation of *L. castroviejoi* and *L. corsicanus* relates to that of the other hare species from Europe, we conducted principal component analyses (PCA) with PLINK 2.00 (Chang et al., 2015), using subsets of polymorphic positions among all species analysed (bi-allelic), separated by a minimum of 50 kb and called in all samples. Additionally, we assessed the genetic structure and potential admixture using ADMIXTURE (Alexander et al., 2009), based on the previous pruned dataset. Multiple runs were performed for varying number of clusters (K) ranging from 2 to 5 (the latter corresponding to the number of included classified species), with 100 bootstrap replicates and 10 cross-validation each. Additional K values (K=6 and 7) were run only for chromosome 20 to reduce computational time. The optimal number of clusters (K) was determined by evaluating cross-validation errors.

We inferred evolutionary relationships and potential migration events among hare species using TreeMix v. 1.13 (Pickrell et al., 2010; Pickrell & Pritchard, 2012). We estimated allele frequencies for the pruned SNPs and ran the TreeMix model with the bootstrap option, with sites grouped in blocks of 500 SNPs to account for linkage disequilibrium. *Lepus americanus* was set as the root. The best tree topology was inferred following the maximum likelihood approach, and up to five migration events were tested.

### Demographic history of divergence between L. castroviejoi and L. corsicanus

We modelled the demographic history of the divergence between *L. castroviejoi* and *L. corsicanus* using the Bayesian method G-PhoCS (Gronau et al., 2011), based on 2,147 intergenic fragments of 1 kb, separated by a minimum distance of 50 kb. We ran three replicates of a model of an ancestral population of size, *N_eAnc_*, that splits in two descendent populations, *L. castroviejoi* and *L. corsicanus* with sizes *N_eCAS_* and *N_eCOR_*, respectively, at time *T*, without gene flow. A second model was run allowing post-split bidirectional gene flow, *m_CAS-COR_* and *m_COR-CAS_*. Each run involved discarding 100,000 generations as burn-in, followed by 1,000,000 MCMC iterations, sampled every 10 iterations. The runs of each model were combined and checked with Tracer v1.7.1 (Rambaut & Drummond, 2014). Scaled demographic parameters obtained from G-PhoCS were converted applying theta=4N_e_*µ, τ=T*µ/g and M=m/µ, where N_e_ is the effective population size (in numbers of individuals), g is the average generation time in years, T is the absolute population divergence time in years, µ is the mutation rate in substitutions/site/generation, and m is the probability of migration in a single generation. We considered a mutation rate of 2.8x10^-9^ substitutions/site/generation (Seixas et al., 2018) and a generation time of two years (Marboutin & Peroux, 1995).

### Detection of introgression

We sought to detect evidence of introgression from other European hare species into either of the two focal sister species using the other as a reference. This could correspond to admixture events occurring after the two sister species split. We first used *D*-statistics (ABBA-BABA tests) (Green et al., 2010; Durand et al., 2011) to compare the number of derived alleles shared by *L. castroviejoi* (P2) and *L. corsicanus* (P1), with each of *L. granatensis*, *L. europaeus* and *L. timidus* (P3), using *L. americanus* as outgroup. Higher introgression into *L. castroviejoi* would result in an excess of shared derived alleles between P2 and P3, relative to P1-P3. For each test, we used a SNP subset with a minimum genotype quality of 10, minimum coverage of 4X (the coverage threshold was relaxed to 3X in the model considering *L. europaeus* as potential donor, given the lower general coverage of these genomes), and a maximum of 60X per SNP. The *D*-statistics were computed using the *Dsuite* toolkit (Malinsky et al., 2021), considering |z-score| ≥ 3 as evidence of gene flow. Additionally, the *f*_4_ statistics (Reich et al., 2009; Pickrell & Pritchard, 2012) were calculated using the fourpop module from the Treemix package (Pickrell et al., 2010; Pickrell & Pritchard, 2012).

To characterize how inferred introgression was distributed along the genome of *L. castroviejoi*, we first estimated the *f*_dM_ statistics (Malinsky et al., 2015; Malinsky et al., 2018) in windows of 25 kb with a minimum of 100 SNPs per window, using Genomics General (https://github.com/simonhmartin/genomics_general). *f*_dM_ values vary from -1 to 1, indicating gene flow between P2 and P3 (>0), or P1 and P3 (<0). The attributions of species to P1, P2 and P3 roles was done as described for the *D-statistics*. *f*_dM_ values were computed using SNPs with at least one valid genotype per population (-- minPopCalls) in at least 75% of the samples (--minCalls).

The analyses described above identified *L. granatensis* as the primary donor of alien genetic variation in *L. castroviejoi* (see Results) and we further characterized this exchange using a statistics based on genetic distances, *d_XY_* (Nei, 1987). Introgression from *L. granatensis* should lead to marked decreases of *d_XY_* between *L. granatensis* and *L. castroviejoi* relative to *L. granatensis*-*L. corsicanus* in affected genomic windows. We estimated *d_XY_* in these species pairs in windows of 25 kb containing at least 250 sites with more than 4X coverage, less than 60X coverage and genotype coverage over 10X. Again, only sites with valid genotypes in at least 75% of the samples (--minCalls), and with at least one valid genotype per species (--minPopCalls) were retained. We identified outlier windows based on a z-score test was performed on the ratio *d_XY_* _(*L. granatensis-L. corsicanus*)_ */ d_XY_* _(*L. granatensis-L. castroviejoi*)_.

Finally, we inferred ancestries along the genome using the ancestry deconvolution method implemented in ELAI (Efficient Local Ancestry Inference) (Guan, 2014). ELAI uses a two-layer hidden Markov model (HMM) based on linkage-disequilibrium (estimated from unphased genomes in our case) to infer variations of ancestry along the genomes of a potentially admixed population (target population hereafter), using other samples as proxies of the parental donor populations. We ran one model using *L. castroviejoi* as target, and four potential ancestries: *L. corsicanus* (to recover the native ancestry), *L. granatensis*, *L. europaeus, L. timidus*, and another model inverting the roles of *L. castroviejoi* and *L. corsicanus*. Assuming that differential introgression events affecting *L. castroviejoi* and *L. corsicanus* occurred after their split, the inferred time of split was used as number of admixture generations. Three independent runs for each model were performed, using 20 Expectation Maximization (EM) steps. The results from the three independent runs were averaged, and the ancestry for each genomic position was assigned considering the ancestry probabilities for each potential donor: >0.7 was considered homozygous, 0.4-0.7 heterozygous, and <0.4 uncertain. For each haplotype, genome tracts were formed by merging consecutive positions with the same ancestry. Given that *L. granatensis* is known to harbour introgressed fragment of *L. timidus* origin (Seixas et al., 2018), we quantified, in *L. castroviejoi*, consecutive tracts of *L. granatensis* and *L. timidus* ancestry (i.e. showing a junction between the allospecific sources), which may be marks of *L. timidus* segments transmitted via and admixed *L. granatensis* (second-hand introgression).

We then estimated the age of introgression events based on average introgression tract length estimated using ELAI, under a model involving a single past admixture event. Tract lengths are informative about the timing of introgression because recombination progressively erodes introgression tracts with time (Pool & Nielsen, 2009; Liang & Nielsen, 2014). Under a simple neutral model, the average introgression tract lengths (*L*) is a function of the proportion of admixture (*f*), the recombination rate in Morgans per base pair (*r*) and the time since the introgression event (*t)*: L = [(1 – *f*) *r* (*t* – 1)]^-1^, from Racimo et al. (2015) (see also Duranton et al., 2020). We also estimated introgression time from *L. timidus* into *L. granatensis* using the data from Seixas et al. (2018) and the formula described above. We used a generation time of 2 years (Marboutin & Peroux, 1995) and the recombination rate inferred in rabbits (r = 1.0 x 10-8; Chantry-Darmon et al., 2006), but to remove the uncertainty of the latter assumption, we focused on the ratios of estimates of introgression times and not on the estimated chronological dates.

### Functional enrichment analysis of introgressed regions

To investigate potential functional impacts of introgression from *L. granatensis* into *L. castroviejoi*, we used g:Profiler (Raudvere et al., 2019) and its g:SCS multiple test correction to conduct a functional enrichment analysis of genes overlapping introgression fragments. The analysis was done considering the union of genes overlapping windows with at least 0.5 introgression frequency from *L. granatensis* estimated with ELAI, with outlier windows of the z-score *d*_XY_ test or with the top 0.01% *f*_dM_ windows (P1 – *L. corsicanus*; P2 – *L. castroviejoi*; P3 – *L. granatensis*), which concern introgressed fragments at high frequency. Also, given that Seixas et al. (2018) inferred that introgression from *L. timidus* into *L. granatensis* was partly promoted by selection, we then tested whether introgression of *L. timidus* fragments into *L. castroviejoi* via *L. granatensis* could have concerned a set of functionally meaningful genes. Functional enrichment analyses were conducted on the set of genes in fragments with *L. granatensis*-*L. timidus* junctions found in *L. castroviejoi*, which are candidates to have been transmitted by the admixed *L. granatensis* population (see Results). For these tests, the background list of genes contained those in all portions of the genome retained for each analysis.

## Results

### The history of divergence of L. castroviejoi and L. corsicanus

Whole genome shotgun resequencing resulted in the data statistics detailed in Table S1. The PCA containing data from all hare species from Europe (*L. castroviejoi*, *L. corsicanus*, *L. granatensis*, *L. europaeus* and *L. timidus*), based on 2,999,121 SNPs, revealed distinct clustering for each species, except for *L. corsicanus* and *L. castroviejoi* which clustered together (Figure 1B, S1). This genetic closeness of the sister species was suggested by ADMIXTURE analysis at K=4 (the best K), where *L. corsicanus* and *L. castroviejoi* remained indistinguishable (Figure S2, S3). Restricting the PCA to *L. corsicanus* and *L. castroviejoi* (based on 433,825 SNPs) did allow separating the species (Figure 1B, S1B).

Treemix confirmed the sister status of *L. castroviejoi* and *L. corsicanus* and suggested that *L. europaeus* is their closest relative, in keeping with Ferreira et al. (2021) (Figure 1C). The inclusion of migration events (m) substantially reduced the residuals of the model for *L. castroviejoi* and *L. corsicanus*. Migration edges involving the ancestor of *L. castroviejoi* and *L. corsicanus* were inferred with *L. timidus* (at m=3), and involving *L. castroviejoi* only with *L. granatensis* (m=4) (Figures 1C and S4).

Modelling the divergence of *L. castroviejoi* and *L. corsicanus* as a simple split from an ancestral population with constant effective population sizes and no post-split gene flow in G-Phocs suggested that they split ∼47 thousand years ago (kya) (95% Highest Posterior Density, HPD, 30.65 kya - 66.49 kya) (Figure 1D; Table S2). Allowing for bidirectional gene flow between the populations after the split suggested no migration and returned similar estimates of N_e_ and time of divergence (Table S2).

### Detection and characterization of introgression

The *D*-statistics revealed significantly stronger signs of introgression (z score > 3) from each of the three potential donor species (P3; *L. granatensis*, *L. timidus*, *L. europaeus*) into *L. castroviejoi* (P2) than into *L. corsicanus* (P1) (Figure 2A). The strongest signal of introgression involved *L. granatensis*, as also supported by the *f*_4_ statistics (Figure 2A, Table S3). Focusing on the contribution of *L. granatensis*, the window-based regression model of *d_XY_* between *L. granatensis* and either *L. corsicanus* or *L. granatensis* explained 96.96% of the variation, and identified 418 outlier windows (z-score > 3 or < -3), of which 396 were indicative of introgression from *L. granatensis* into *L. castroviejoi* (Figure 2B). These results were also confirmed by the *f*_dM_ scans (Figure 2C, S5).

**Figure 2:**
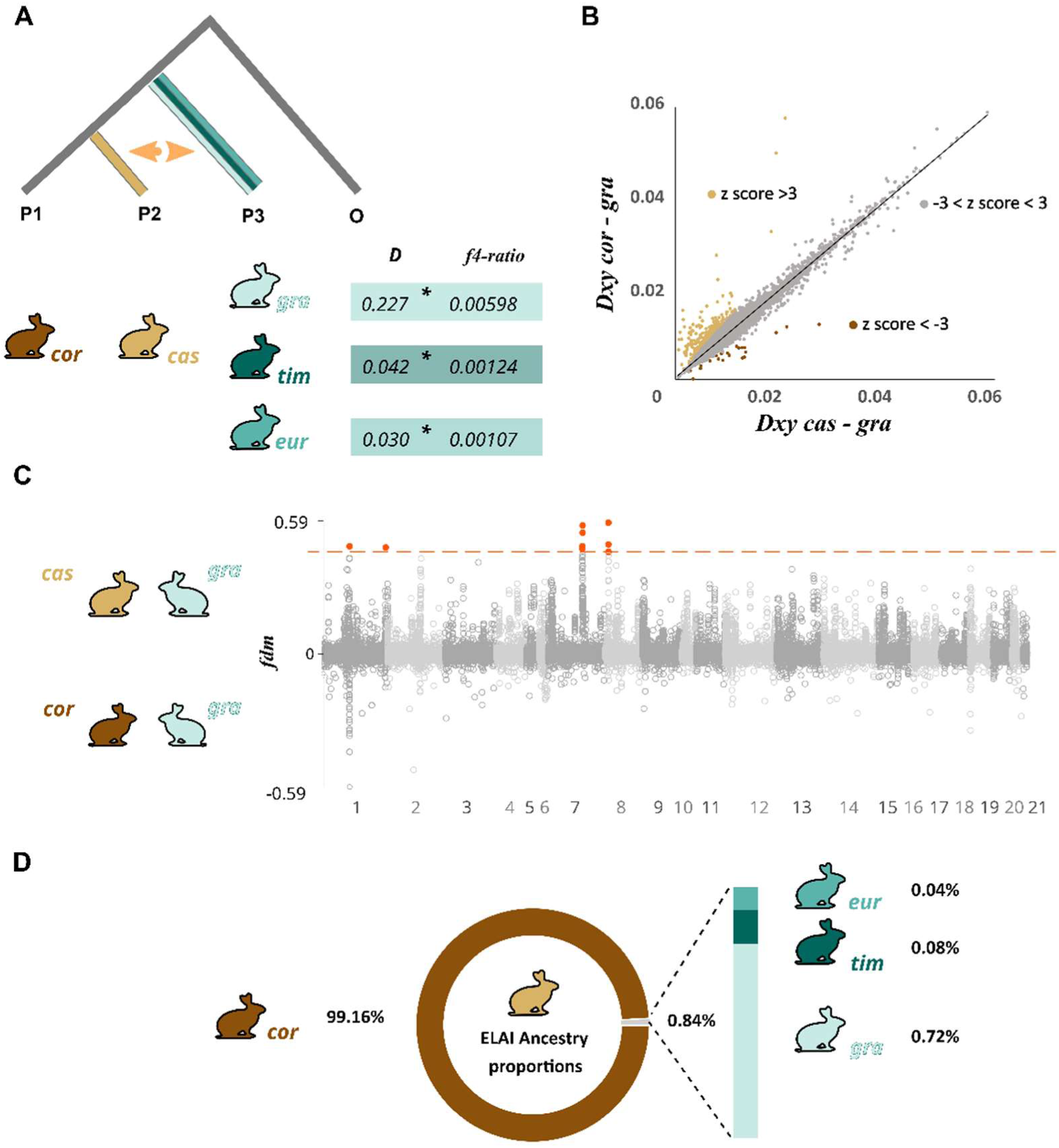
Evidence of admixture. **A.** *D*-statistics tests of introgression into the sister *L. castroviejoi* (*cas*; P2) or *L. corsicanus* (*cor*; P1) from three potential donor species, *L. granatensis* (*gra*, P3), *L. timidus* (*tim*, P3) *L. europaeus* (*eur*, P3). *L. americanus* was used as outgroup (O). **B.** Correlation of genetic distances d_XY_ between *cas-gra* and *cor-gra* in windows of 25 kb along the genome. Z-score outliers are highlighted. **C.** *fdm* tests of admixture from *gra* into *cas* or *cor* along the genome in windows of 25 kb. Orange dots indicate top 0.01% values. **D.** Proportions of genomic ancestry of the *cas* genome from *cor* (identifying native ancestry), *gra*, *tim* and *eur* inferred using ELAI.

Using the ELAI local ancestry inference method, placing *L. castroviejoi* as the target species for admixture assigned 99.16% of its genomes to a *L. corsicanus-*like parent, while only 0.74% was assigned to other ancestries: 0.72% to *L. granatensis*, 0.08% to *L. timidus* and 0.04% to *L. europaeus* (Figure 2D, Table S4, S5). Most of the inferred contributions of *L. granatensis* to *L. castroviejoi* were found at low frequencies, but ∼5% of these tracts were detected at frequencies ≥0.5 (0.58% were found fixed across our samples) (Figure S6). In contrast, using *L. corsicanus* as the target population for admixture, 99.88% of the genomes was assigned to *L. castroviejoi* (again, indicative of native ancestry), with just 0.075% being assigned to other ancestries (Table S8). Ninety tracts of allospecific ancestry in *L. castroviejoi* were found to harbour *L. granatensis*-*L. timidus* junctions, which can mark introgression of *L. timidus* via the admixed *L. granatensis*.

To assess the consistency of introgression evidences inferred across the different methods used here, we calculated *d_XY_* and *f*_dM_ values only for regions detected as introgressed with ELAI, binned by introgression frequency. For ELAI introgression tracts from *L. granatensis* into *L. castroviejoi*, *f*_dM_ values were strongly positively correlated with ELAI-inferred introgression frequencies, while *d_XY_* between *L. castroviejoi* and *L. granatensis* was negatively correlated with frequencies (Figure S7). Focusing on the 396 *d_XY_* z-score outliers, we found that they were included in the top 0.01% *f*_dM_ windows (from 80,010 windows), and in 267 of them ELAI also detected introgression. These results suggested an overall consistency of local genomic inferences of introgression across the different approaches.

Using the mean of ancestry tract lengths and admixture proportions, we estimate that introgression from *L. timidus* into *L. castroviejoi* (tim2cas) is 1.7 times older than that from *L. granatensis* into *L. castroviejoi* (gra2cas), while introgression from *L. europaeus* into *L. castroviejoi* occurred in an intermediate time frame (1.2 times older than gra2cas) (Table S6, S7). We also calculated relative introgression dates, in comparison with patterns of *L. timidus* to *L. granatensis* introgression (tim2gra), using the data from Seixas et al. (2018). We estimate that tim2cas introgression was 1.3 times older than tim2gra, and that tim2gra was 1.4 times older than gra2cas. We also did the calculations separately for the distal and proximal chromosomal regions, which have been shown to have different recombination rates in hares (Seixas et al., 2018), and obtained similar results (Table S7). To summarize, we estimate that introgression ages rank as follows (from ancient to recent): tim2cas > tim2gra > gra2cas.

### Functional enrichment of introgressed genes

Among the collection of genes overlapping tracts with high introgression frequency of *L. granatensis* origin in *L. castroviejoi*, no enrichment of a particular ontology term was found. Yet, several genes in these regions are linked with cell metabolism (biological regulation, regulation of biological process, regulation of cellular process, cell communication, signalling, signal transduction, voltage-gated sodium channel complex) (Figure S8, Table S8).

The functional enrichment analysis of the 90 genes overlapping tracts with *L. granatensis-L. timidus* junctions in *L. castroviejoi* detected enrichment of olfactory receptor activity genes, and also several genes linked to other signalling activities, such as signalling receptor activity, G protein-coupled receptor activity, transmembrane signalling receptor activity and molecular transducer activity (Figure S8).

## Discussion

Taking advantage of the vicariance of the two sister species, *L. castroviejoi* and *L. corsicanus*, we reveal the complex history of introgression affecting *L. castroviejoi* from three other species in Western Europe. Though with a reduced fraction of the genome affected by introgression, *L. castroviejoi* today is a mosaic of multiple ancestors that witnesses a history of multiple contacts and hybridization events.

*The divergence between* L. castroviejoi *and* L. corsicanus *predates range revolutions leading to multi-way admixture in hares* We analysed whole genome data to trace the history of divergence between the sister species *L. castroviejoi* and *L. corsicanus*, in the context of the genetic diversity currently present across European hare species. The genetic similarity we found between these species, which are poorly separated in analyses that consider the differentiation among all European species (Figure 1B), confirms previous results based on a few markers (Alves et al., 2008a; Melo-Ferreira et al., 2012) but also on whole exome analysis of single individuals (Ferreira et al., 2021). We estimated the divergence between *L. castroviejoi* and *L. corsicanus* to have occurred at Late Pleistocene, ∼47 kya (Figure 1D).

All our estimates show that the introgression fraction is much higher in *L. castroviejoi* than in *L. corsicanus*. For example, the foreign genetic contribution based on ELAI in *L. castroviejoi* (Figure 2D, Table S3) was ten-fold higher than in *L. corsicanus* (0.74% and 0.075% respectively; Table S4). Yet the *L. granatensis* contribution to the genome of *L. corsicanus* was unexpected since they are allopatric. The very low alien contribution in *L. corsicanus* could result from artefacts of the methods (false positives), but we must also consider that our sample of *L. corsicanus* comes from an island (Corsica) where *L. europaeus* and *L. granatensis* are reported to have been introduced (Pietri et al., 2011).

Mitochondrial DNA in *L. castroviejoi* and *L. corsicanus* has been shown to be entirely of *L. timidus* origin (Melo-Ferreira et al., 2012), which suggests the ancestor of the two species hybridized with *L. timidus* and captured its mitochondrial genome. Our Treemix analysis with three migration edges also suggested this route of introgression, which may witness these ancient introgression events (Figure S4) that must have occurred at least 47 kya, the estimated divergence time of the sister species. If the *L. timidus* nuclear introgression resulted solely from this ancient event, the average size of the introgression fragments should be around 4 kb, but our estimation rather points to an average around 25 kb (with a median ∼18 kb). The probability to observe a tract longer than 18 kb is about 1.3% (exp(-18/4)) (following Racimo et al., 2015), but our results show that half of them are longer, suggesting that *L. castroviejoi* was affected by a second wave of more recent introgression from *L. timidus*.

We note however that the methods we used would not have been able to detect high frequency introgression from *L. timidus* into both *L. castroviejoi* and *L. corsicanus* in the same genomic regions. With our design, we can only detect introgression when it is highly asymmetric between the two sister species. Nevertheless, the direction of the asymmetry we find and the hardly detectable introgression into *L. corsicanus* indicates that most of the detectable introgression occurred in the Iberian Peninsula, where *L. castroviejoi* is present together with two other species also affected by more recent introgression from *L. timidus*.

### Post-glacial hybridization affected the gene pool of L. castroviejoi in the Iberia Peninsula

*L. castroviejoi* is shown here to be a mosaic of at least three other ancestries (Figure 2; Table S4). Such multiway admixture pattern is not uncommon in nature (e.g. Fontaine et al., 2015; Toews et al., 2018; Grant & Grant, 2020; Martin et al., 2020). The overall foreign contribution to *L. castroviejoi* is less than 1% (Figure 2D), which explains the lack of detection of admixture in a recent microsatellite study (Costa et al., 2024). The strongest foreign contributor to *L. castroviejoi* is *L. granatensis*, with estimates of introgression affecting 0.465 to 0.895% of the individual genomes analysed (Table S4). Our results must be reconciled with what is known about introgression into the other species of the Iberian Peninsula, and *L. granatensis* has been studied in greatest detail in that perspective (Alves et al., 2008b; Melo-Ferreira et al., 2012; Melo-Ferreira et al., 2014; Seixas, 2017; Seixas et al., 2018). This species is endemic to the Peninsula, which it has almost entirely recolonised from a Southern refugium after the last glacial period (Marques et al., 2017; Seixas et al., 2018). Introgression from *L. timidus* also affected *L. granatensis*, along a south-north gradient (about 1.3% of its genome in the south to about 2.4% in the north), and was estimated to have occurred ∼7 kya (Seixas et al., 2018).

Using average tract lengths and fractions of introgression, we ordered the estimated times of the different admixture events by estimating their ratios. Considering that the different introgression events are independent, we estimated that introgression followed this order, from oldest to most recent: tim2cas > tim2gra > gra2cas (Table S7). According to these inferences, *L. castroviejoi* would have hybridized with *L. granatensis* that was already introgressed by *L. timidus*. We must thus consider the possibility that introgression of *L. timidus* into *L. castroviejoi* occurred via a *L. granatensis* intermediate (second-hand introgression scenario, which we will summarize by gratim2cas). This could account for our inferences of *granatensis*-*timidus* junctions in the *castroviejoi* genomes, an occurrence that would be rather unlikely given the sparsity of introgression tracts, if the two introgression events were independent. Our recalculation of the dates of introgression into *L. castroviejoi* when considering *L. granatensis* and *L. timidus* tracts together, and merging adjacent *L. granatensis* and *L. timidus* tracts, is compatible with the second-hand introgression hypothesis, since the gratim2cas time is estimated more recent than tim2gra (Table S7).

We must, however, be cautious about these admixture time estimates given several possible sources of bias. By estimating time ratios, we attempted to eliminate one nuisance parameter, recombination rate. Doing so makes the assumption that recombination rate does not vary significantly among species (which we cannot verify), but recombination rate is known to vary along the genome and we averaged data across the whole genome. However, separate estimates based on partition among broad-scale higher and lower recombination genomic regions (chromosome ends versus centres) were concordant with the global analysis. Yet, the decreased power to detect small introgression tracts (Seixas et al., 2018) may have led to underestimates of the times of the events, and selection may also cause variations of admixture rates along the genome. Our approach supposes that similar biases affected all comparisons, and is thus dependent on this assumption.

*L. europaeus* appears as the minor contributor to the genome of *L. castroviejoi*, with a small number of tracts. We estimated introgression to have occurred in roughly the same timeframe as the other such events, likely when *L. europaeus* expanded to western Europe (Stamatis et al., 2009). Overall, our results broadly point to a short timeframe, starting at early Holocene, when multi-way admixture among several species occurred in the Iberian Peninsula.

## Conclusion

We have found that, despite complete fixation of mitochondrial genomes of *L. timidus* origin in both *L. castroviejoi* and *L. corsicanus*, suggesting ancient introgression before the divergence of these sister species, *L. castroviejoi* has been more affected than its sister by nuclear introgression from this origin. This *L. castroviejoi*-specific *timidus* nuclear introgression must have occurred in the Iberian Peninsula, from which *L. castroviejoi* is endemic, and where *L. timidus* is known to have been present in the past, as attested by its contribution to the genome of another Iberian endemic*, L. granatensis* (Seixas et al., 2018). Our results suggest that the (modest) *L. timidus* nuclear contribution to the *L. castroviejoi* genome may have occurred through a *L. granatensis* intermediate.

We found that the segments carrying *L. timidus*-*L. granatensis* junctions, and thus potentially transmitted through *L. granatensis*, contain a set of genes with functions related to cell signalling and an enrichment in olfactory receptor activity functions. Variants for such functions could participate to adaptation in a given region or be universally adaptive, irrespective of the host species. Finally, our work shows that footprints of past hybridization events can convey indirect information to reconstruct complex past biogeographic histories.

## Author Contributions

J.M.-F., J.P.M., P.B. coordinated the study; J.M.-F., J.P.M., P.B., P.C.A. designed research; J.M.-F. acquired funding; F.B., C.P. contributed with samples; L.F. performed laboratory work; J.S., J.P.M. analysed data; J.S., J.P.M., J.C., J.Q., P.C.A., P.B., J.M.-F. contributed to the interpretation of the results; J.S., J.P.M., J.M.-F. wrote the paper; J.S., J.P.M., L.F., J.C., J.Q., C.P., P.C.A., P.B., J.M.-F. revised the paper and approved the final version.

## Acknowledgements

This work is dedicated to the memory of Fernando Ballesteros. All authors agree with his inclusion as co-author and the corresponding author accepts all responsibility for the co-author’s data and contribution.

## Funding

This work was funded by project HybridChange (PTDC/BIA-EVL/1307/2020), supported by Portuguese national funds through Fundação para a Ciência e a Tecnologia, FCT. JMF was supported by FCT (CEEC contract with reference 2021.00150.CEECIND). JC was supported by an FCT PhD grant (reference UI/BD/154450/2022). Additional support was obtained from the European Union’s Horizon 2020 Research and Innovation Programme under the Grant Agreement Number 857251.

## Conflict of Interest Statement

The authors declare no conflict of interest.

## Data Availability Statement

All data will be deposited in NCBI’s SRA. The generated pseudo-reference genome will be available at Dryad. Custom scripts will be available at GitHub.

## Notes

### Competing Interest Statement

The authors have declared no competing interest.

