## Supplementary Material for "Genome admixture among four hare species in Iberia: focus on the broom hare (*Lepus castroviejoi*)"

**Supplementary Figures:**

**Supplementary Tables:**

Table S1 ………………………………………………………………………………….. 10

Table S2 ………………………………………………………………………………….. 13

Table S3 ………………………………………………………………………………….. 14

Table S4 ………………………………………………………………………………….. 14

Table S5 ………………………………………………………………………………….. 15

Table S6 ………………………………………………………………………………….. 16

Table S7 ………………………………………………………………………………….. 17

Table S8 ………………………………………………………………………………….. 18

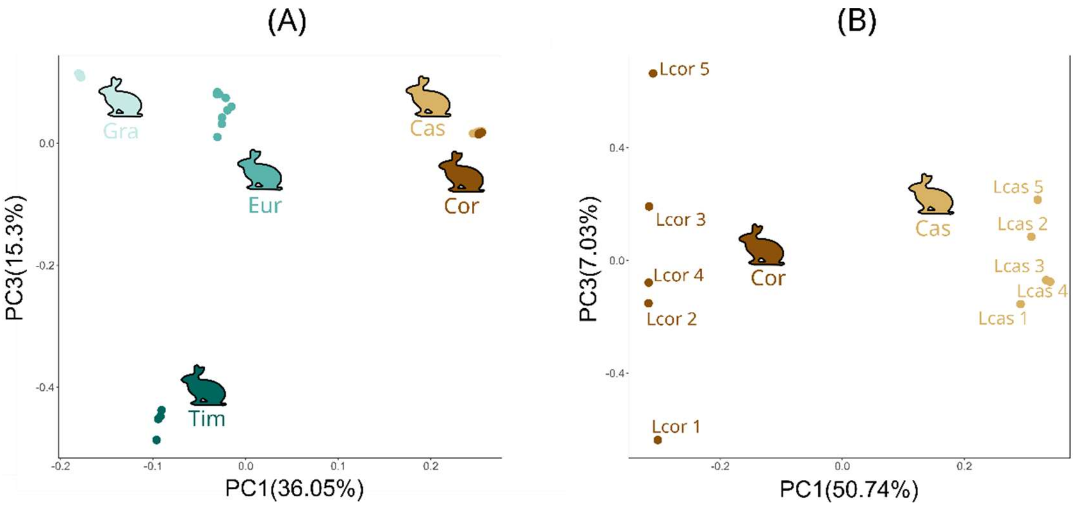

**Figure S1.** Principal Component Analysis (PCA) based on (A) SNP data of European hare species (2,999,121 SNPs) and (B) for *L. castroviejoi* and *L. corsicanus* (433,825 SNPs). Principal components 1 and 3 are shown with the proportion of variance explained by each axis. *Cas – L. castroviejoi*; *Cor – L. corsicanus*; *Gra – L. granatensis*; *Eur – L. europaeus*; *Tim – L. timidus*.

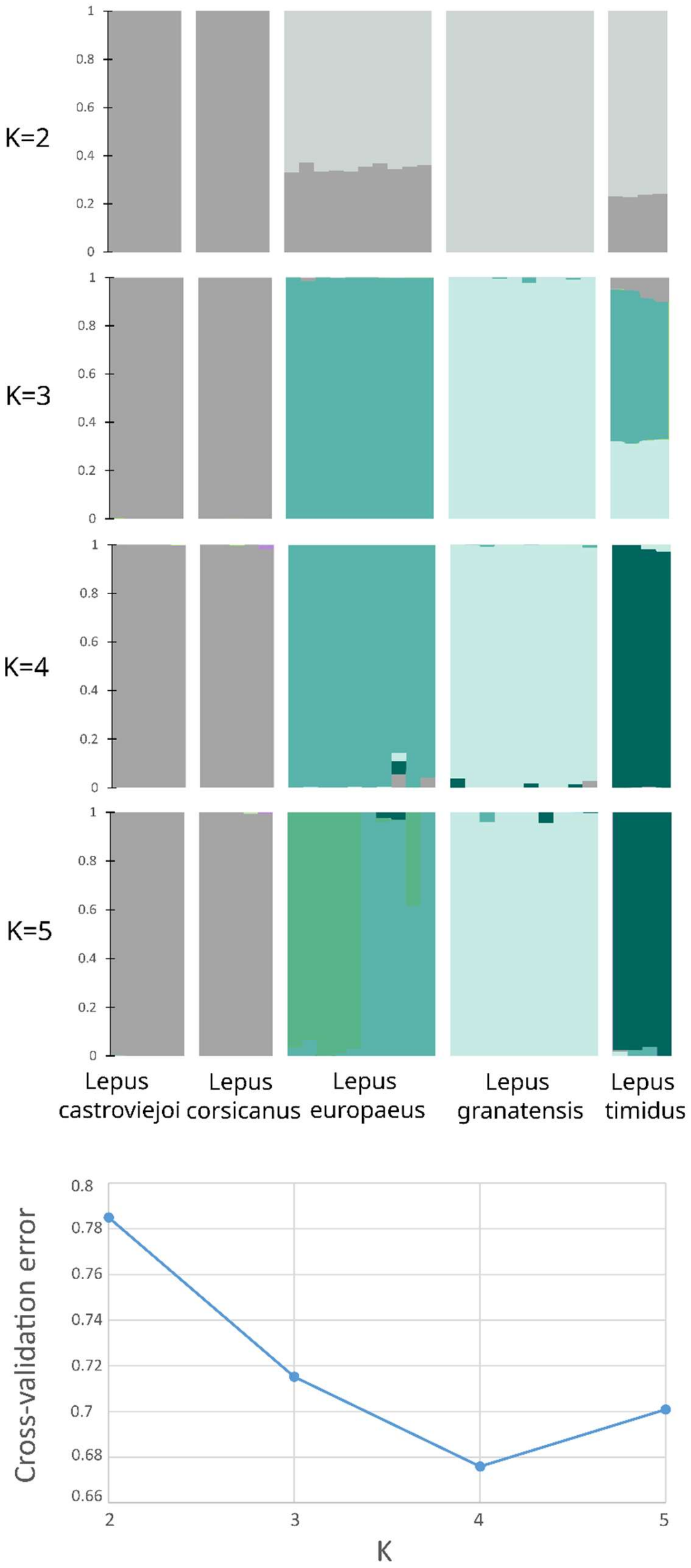

**Figure S1.** ADMIXTURE analysis (top panel; based on ~3 million SNPs filtered by linkage disequilibrium) for European hare species, considering from K= 2 to K=5 number of populations. The cross-validation errors are shown at the bottom.

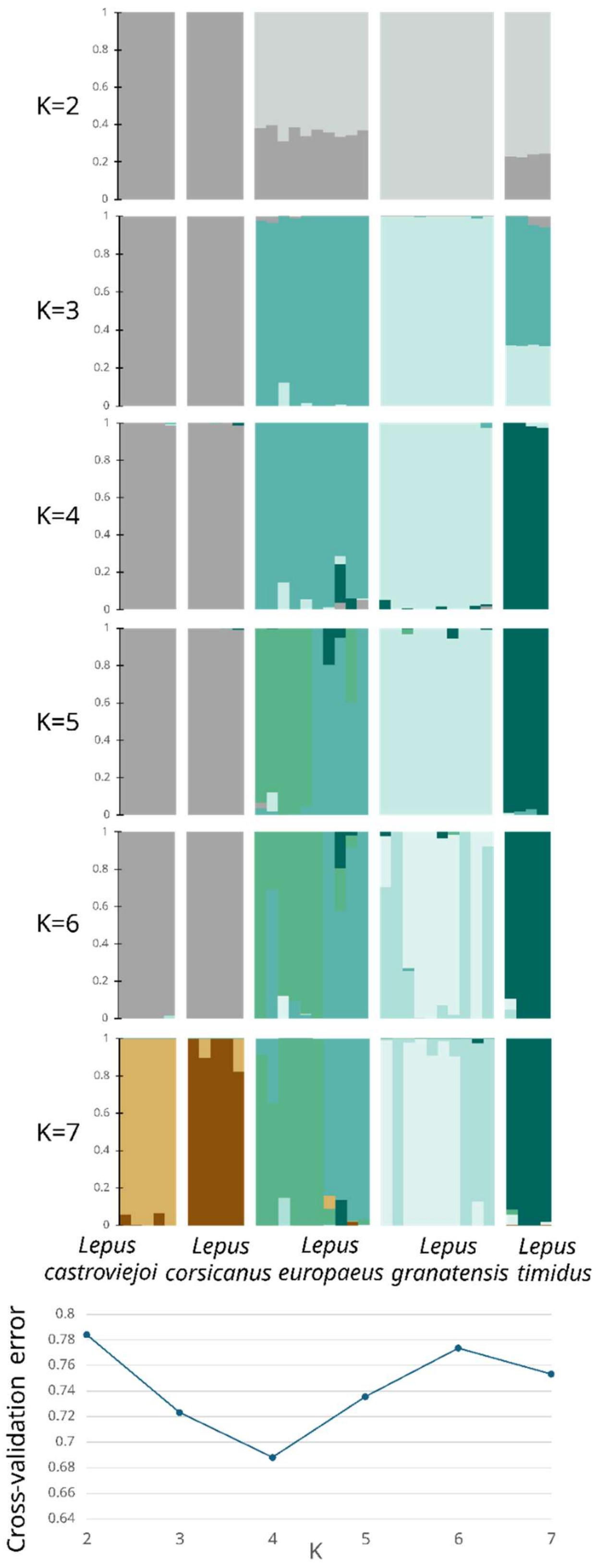

**Figure S2.** ADMIXTURE analysis (top panel) for European hare species, considering K=2 to K=7 number of populations, based on independent SNPs on chromosome 20. The cross-validation errors are shown at the bottom.

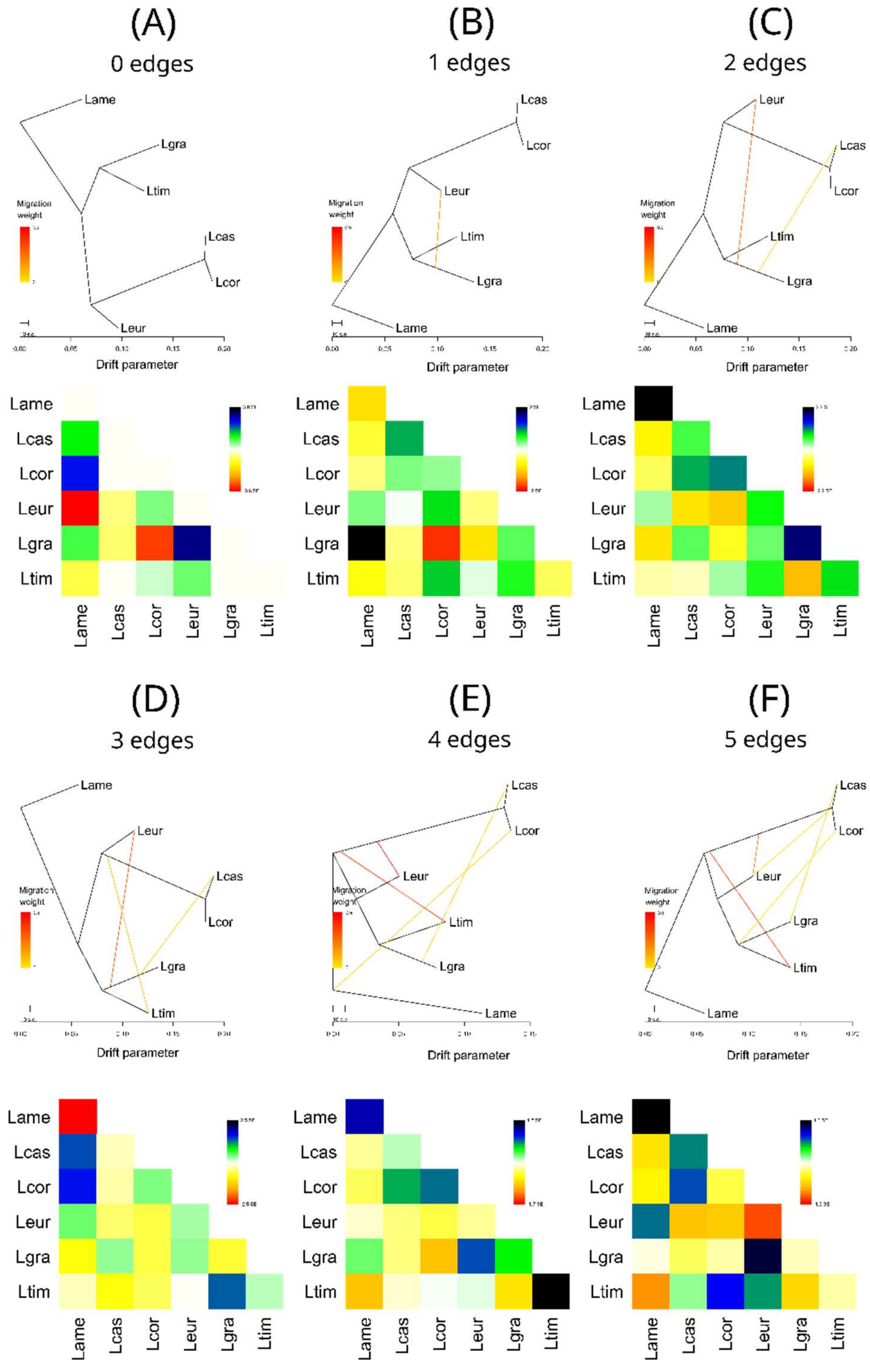

**Figure S3.** Relationships among species of hares from Europe inferred using Treemix and *L. americanus* as the outgroup. model without migration (A), and with 1 (B), 2 (C), 3 (D), 4 (E), and 5 (F) migration edges. Model residuals are shown at the bottom of each topology. Lame – *Lepus americanus*; Lcas – *Lepus castroviejoi*; Lcor – *Lepus corsicanus*; Leur – *Lepus europaeus*; Lgra – *L. granatensis*; Ltim – *Lepus timidus*.

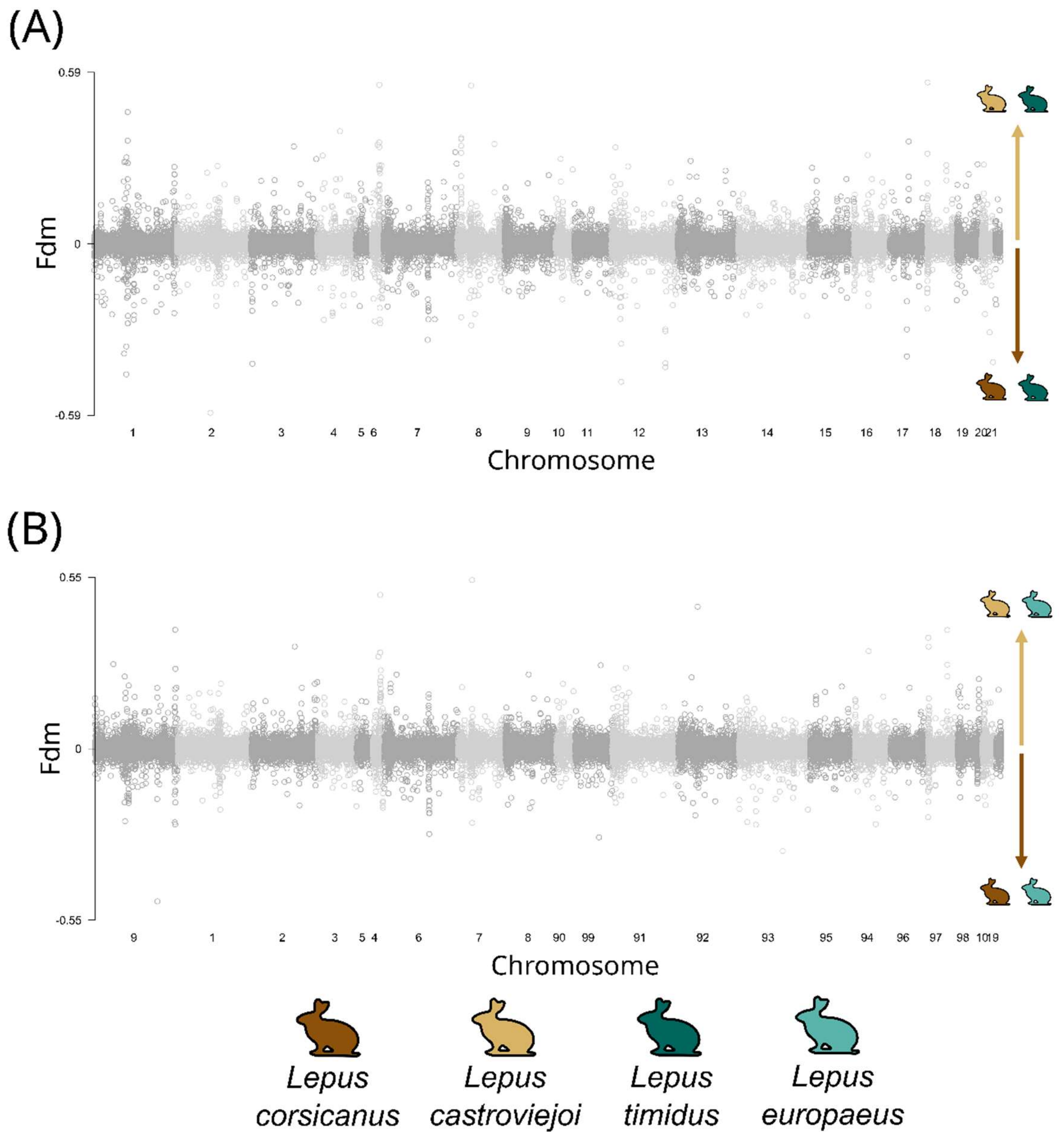

**Figure S5.** Genome wide $f_{dM}$ values for a model where P1 – *L. corsicanus*, P2 – *L. castroviejoi*, O – *L. americanus*, and two P3 tested: (A) P3 – *L. timidus* and (B) *L. europaeus*. Positive values suggest geneflow between P2 and P3, while negative values indicate gene flow between P1 and P3.

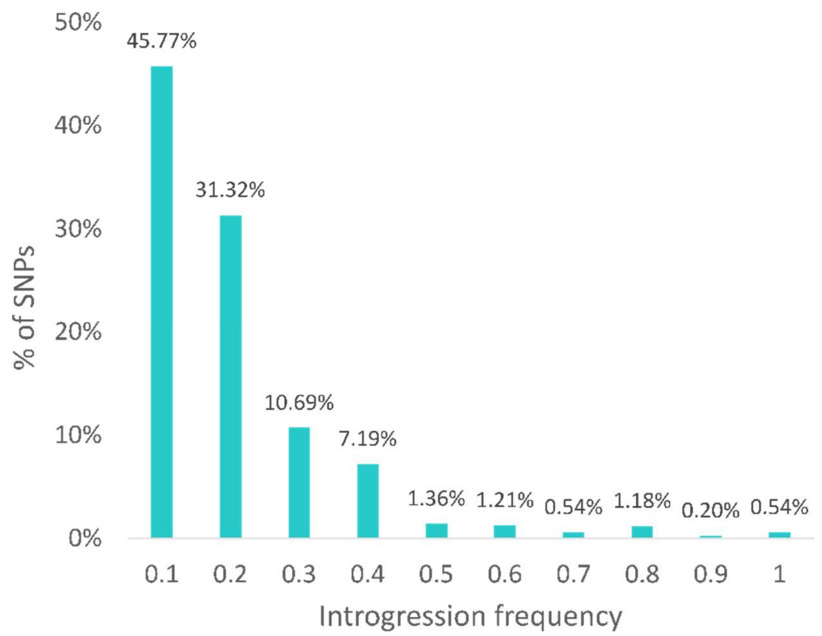

**Figure S6.** Distribution of the frequencies of introgression of SNPs with *L. granatensis* ancestry inferred in the *L. castroviejoi* genome using ELAI.

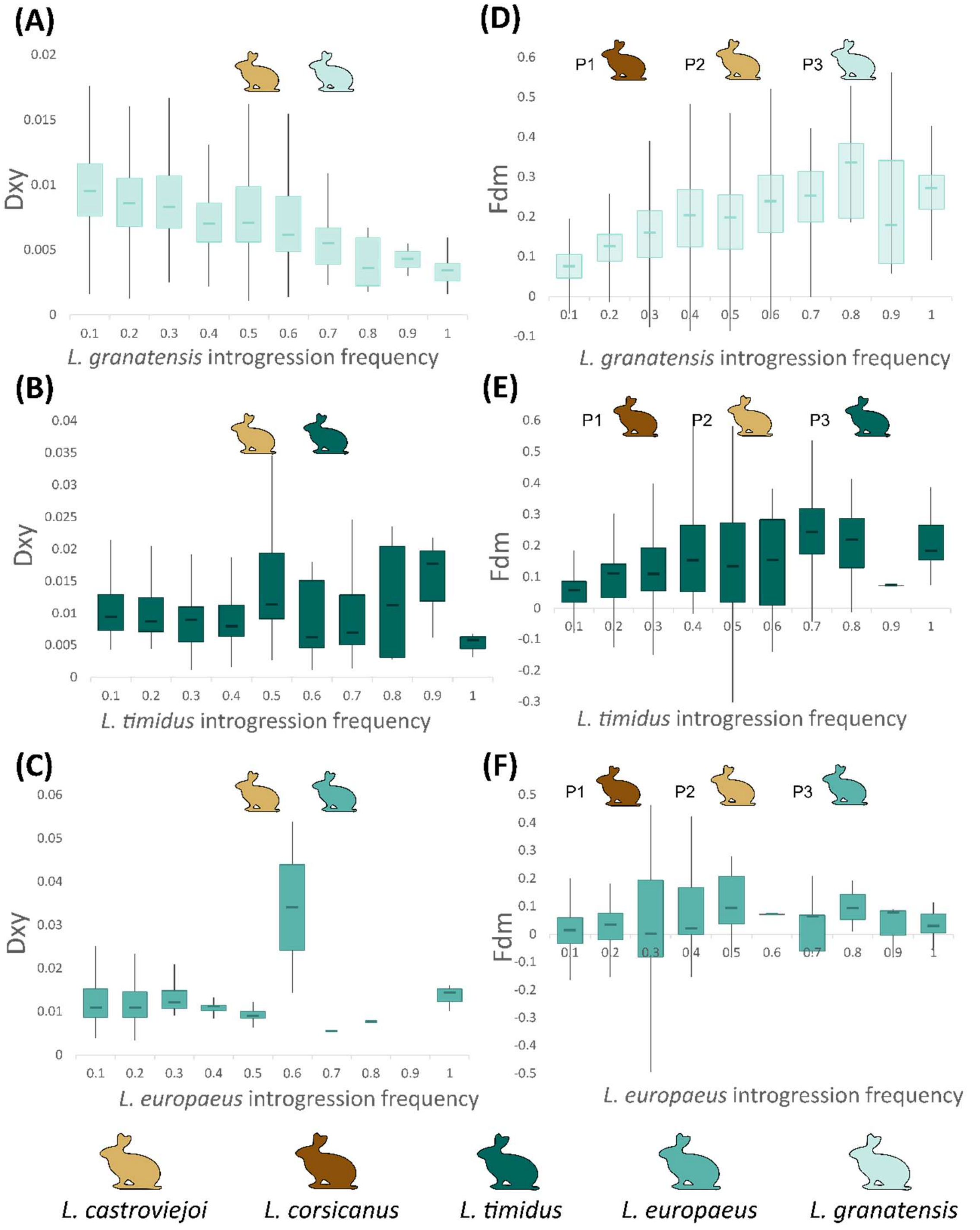

**Figure S7.** Boxplots with $d_{XY}$ estimates per ELAI introgression frequency in *L. castroviejoi* from different origins. (A) *L. castroviejoi-L. granatensis*; (B) *L. castroviejoi-L. timidus*; (C) *L. europaeus-L. castroviejoi*; and boxplots of $f_{DM}$ values (P1 – *L. corsicanus*; P2 – *L. castroviejoi*) per ELAI introgression frequency in *L. castroviejoi* using as P3 (D) *L. granatensis*; (E) *L. timidus*; and (F) *L. europaeus*, and *L. americanus* as outgroup.

(A)

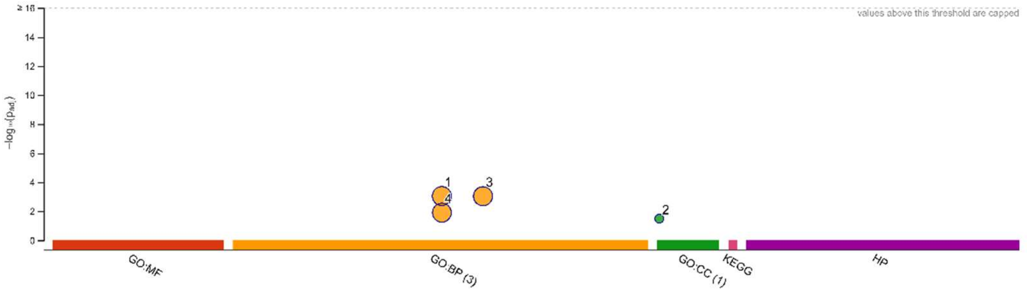

| ID | Source | Term ID | Term Name | $p_{adj}$ (query_1) |
|---|---|---|---|---|
| 1 | GO:BP | GO:0050789 | regulation of biological process | $8.921\times10^{-4}$ |
| 2 | GO:CC | GO:0001518 | voltage-gated sodium channel complex | $3.216\times10^{-2}$ |
| 3 | GO:BP | GO:0065007 | biological regulation | $9.183\times10^{-4}$ |
| 4 | GO:BP | GO:0050794 | regulation of cellular process | $1.247\times10^{-2}$ |

(B)

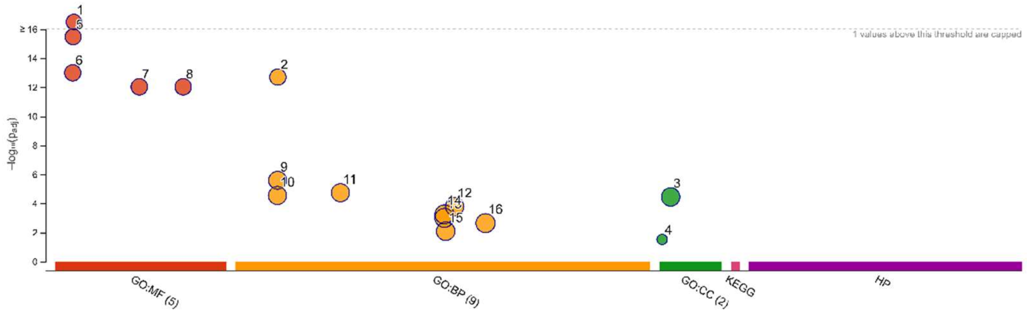

| ID | Source | Term ID | Term Name | $p_{adj}$ (query_1) ↑ |
|---|---|---|---|---|
| 1 | GO:MF | GO:0004984 | olfactory receptor activity | $8.618\times10^{-20}$ |
| 5 | GO:MF | GO:0004930 | G protein-coupled receptor activity | $3.351\times10^{-16}$ |
| 6 | GO:MF | GO:0004888 | transmembrane signaling receptor activity | $1.012\times10^{-13}$ |
| 2 | GO:BP | GO:0007186 | G protein-coupled receptor signaling pathway | $2.008\times10^{-13}$ |
| 7 | GO:MF | GO:0038023 | signaling receptor activity | $9.280\times10^{-13}$ |
| 8 | GO:MF | GO:0060089 | molecular transducer activity | $9.280\times10^{-13}$ |
| 9 | GO:BP | GO:0007165 | signal transduction | $2.561\times10^{-6}$ |
| 11 | GO:BP | GO:0023052 | signaling | $1.834\times10^{-5}$ |
| 10 | GO:BP | GO:0007154 | cell communication | $2.854\times10^{-5}$ |
| 3 | GO:CC | GO:0016020 | membrane | $3.585\times10^{-5}$ |
| 12 | GO:BP | GO:0051716 | cellular response to stimulus | $1.641\times10^{-4}$ |
| 14 | GO:BP | GO:0050794 | regulation of cellular process | $5.642\times10^{-4}$ |
| 13 | GO:BP | GO:0050789 | regulation of biological process | $9.860\times10^{-4}$ |
| 16 | GO:BP | GO:0065007 | biological regulation | $2.275\times10^{-3}$ |
| 15 | GO:BP | GO:0050896 | response to stimulus | $8.197\times10^{-3}$ |
| 4 | GO:CC | GO:0001917 | photoreceptor inner segment | $3.021\times10^{-2}$ |

**Figure S8.** g:Profiler Gene Ontology analysis genes in genomic regions of *L. castroviejoi* with (A) high introgression frequency of *L. granatensis* origin ($d_{XY}$ z-score outlier windows or showing *L. granatensis* introgression frequencies of at least 50% inferred in ELAI); and with (B) gratim2cas segments (segments of ancestry junctions between *L. granatensis* and *L. timidus* found in *L. castroviejoi*. P-values are shown.

**Table S1.** Samples and whole genome sequencing datasets used in this work. Data were either newly generated or retrieved from previous works ((1) Giska et al. (2019); (2) Seixas et al. (2018); (3) Seixas (2017); (4) Carneiro et al. (2014)). GQ – genotype quality.

| ID | Species | Location | Reference | Mean Depth | Mean GQ | SRA Accession Nr. |
|---|---|---|---|---|---|---|
| **Lcas1** | *L. castroviejoi* | Cantabria, Spain | this work | 14.198 | 39.555 | ### |
| **Lcas2** | *L. castroviejoi* | Alto Sil, León, Spain | (1) | 8.232 | 23.535 | ### |
| **Lcas3** | *L. castroviejoi* | León, Spain | this work | 6.889 | 21.080 | ### |
| **Lcas4** | *L. castroviejoi* | Riano, León, Spain | this work | 6.035 | 18.391 | ### |
| **Lcas5** | *L. castroviejoi* | Cantabria, Spain | this work | 7.115 | 20.556 | ### |
| **Lcor1** | *L. corsicanus* | Corsica, France | this work | 17.897 | 48.700 | ### |
| **Lcor2** | *L. corsicanus* | Corsica, France | this work | 8.910 | 25.496 | ### |
| **Lcor3** | *L. corsicanus* | Corsica, France | this work | 9.594 | 27.180 | ### |
| **Lcor4** | *L. corsicanus* | Corsica, France | this work | 7.472 | 20.214 | ### |
| **Lcor5** | *L. corsicanus* | Corsica, France | this work | 9.248 | 25.815 | ### |
| **Lgra1** | *L. granatensis* | Alcoutim, Portugal | (2) | 18.490 | 53.804 | ### |
| **Lgra2** | *L. granatensis* | Peñaflor, Sevilla,Spain | (2) | 17.596 | 52.610 | ### |
| **Lgra3** | *L. granatensis* | Pancas, Portugal | (2) | 14.980 | 43.422 | ### |

| | | | | | | |
|---|---|---|---|---|---|---|
| **Lgra4** | *L. granatensis* | Idanha, Castelo Branco, Portugal | (2) | 18.692 | 53.866 | ### |
| **Lgra5** | *L. granatensis* | Miguelturra, Ciudad Real, Spain | (2) | 18.884 | 55.586 | ### |
| **Lgra6** | *L. granatensis* | Valpaços, Portugal | (2) | 19.720 | 57.000 | ### |
| **Lgra7** | *L. granatensis* | Algete, Madrid, Spain | (2) | 19.817 | 60.223 | ### |
| **Lgra8** | *L. granatensis* | Valencia province, Spain | (2) | 15.312 | 45.963 | ### |
| **Lgra9** | *L. granatensis* | Sauguillo, Soria, Spain | (2) | 14.660 | 43.491 | ### |
| **Lgra10** | *L. granatensis* | Fontellas, Navarra, Spain | (2) | 20.123 | 58.251 | ### |
| **Leur1** | *L. europaeus* | Cantabria, Spain | (3) | 7.032 | 22.863 | ### |
| **Leur2** | *L. europaeus* | Jaca, Spain | (3) | 8.201 | 27.097 | ### |
| **Leur3** | *L. europaeus* | Villarcayo, Spain | (3) | 9.065 | 28.803 | ### |
| **Leur4** | *L. europaeus* | Álava, Spain | (3) | 7.835 | 26.189 | ### |
| **Leur5** | *L. europaeus* | Navarra, Spain | (3) | 11.083 | 34.632 | ### |
| **Leur6** | *L. europaeus* | Pyrenees, France | (3) | 7.514 | 20.063 | ### |
| **Leur7** | *L. europaeus* | Ukraine | (3) | 10.867 | 26.576 | ### |
| **Leur8** | *L. europaeus* | Germany | (3) | 10.715 | 25.678 | ### |
| **Leur9** | *L. europaeus* | Vienna, Austria | (3) | 9.269 | 20.155 | ### |

| | | | | | | |
|---|---|---|---|---|---|---|
| **Leur10** | *L. europaeus* | Clermont-Ferrand, France | (3) | 5.542 | 15.713 | ### |
| **Ltim1** | *L. timidus* | Borris-in-Ossory, Ireland | (2) | 22.591 | 67.264 | ### |
| **Ltim2** | *L. timidus* | Finland | (2) | 16.746 | 51.877 | ### |
| **Ltim3** | *L. timidus* | Calfreisen, Egga, Switzerland | (2) | 18.550 | 57.349 | ### |
| **Ltim4** | *L. timidus* | Nancy-sur-Cluses, France | (2) | 21.091 | 63.552 | ### |
| **Lame** | *L. americanus* | Lake Inez, Missoula, Montana, USA | (4) | 27.384 | 75.270 | ### |

**Table S2.** Demographic parameters inferred with G-PhoCS for the history of divergence between *L. castroviejoi* (cas) and *L. corsicanus* (cor) for models with and without post-split gene flow (A). Conversions of raw estimates were done by using a generation time of two years and mutation rate μ = 2.8 x 10-9 substitutions/site/generation. Mean values of estimated parameters are presented with 95% HPD intervals in parentheses.

| **G-PhoCS parameter** | **Demographic parameter (95% HPD interval)** | |
|---|---|---|
| | **Model without gene flow** | **Model with gene flow** |
| **theta lcor** | 18 232 (14 688 - 21 876) diploid individuals | 17 304 (13 607 – 21 234) diploid individuals |
| **theta lcas** | 15 571 (12 473 – 18 748) diploid individuals | 14 705 (11 558 – 17 989) diploid individuals |
| **theta root** | 158 143 (122 179 – 218 063) diploid individuals | 158 759 (139 022 – 178 671) diploid individuals |
| **tau root** | 47 000 (30 650 – 66 493) generations | 43 214 (35 543 – 50 986) generations |
| **m lcas > lcor** | - | 0 migrants/generation |
| **m lcor > lcas** | - | 0 migrants/generation |

**Table S3.** $f_4$ statistics based on topology (A,B)(C,D). Negative values indicate gene flow between C and B or between D and A. Positive values imply gene flow between A and C or between B and D. Lame – *L. americanus*; Lgra – *L. granatensis*; Lcas – *L. castroviejoi*; Lcor – *L. corsicanus*; Ltim – *L. timidus*; Leur – *L. europaeus*.

| | | | $f_4$ | | | |
|---|---|---|---|---|---|---|
| **Pop A** | **Pop B** | **Pop C** | **Pop D** | **$f_4$ statistics** | **S.E.** | **Z-score** |
| Lame | Lgra | Lcas | Lcor | -0.00112 | 7.86E-05 | -14.29 |
| Ltim | Lgra | Lcas | Lcor | -0.00076 | 6.07E-05 | -12.51 |
| Lame | Leur | Lcas | Lcor | -0.00012 | 4.25E-05 | -2.75 |
| Ltim | Leur | Lcas | Lcor | 0.00025 | 5.06E-05 | 4.89 |
| Ltim | Lame | Lcas | Lcor | 0.00037 | 5.14E-05 | 7.10 |

**Table S4.** Proportions of the *L. castroviejoi* genome attributed to 4 different ancestries estimated with ELAI, in each analysed individual and overall.

| **Ind** | ***L. corsicanus*** | ***L. granatensis*** | ***L. europaeus*** | ***L. timidus*** |
|---|---|---|---|---|
| **Lcas1** | 98.823% | 1.020% | 0.047% | 0.110% |
| **Lcas2** | 99.097% | 0.805% | 0.028% | 0.070% |
| **Lcas3** | 99.343% | 0.547% | 0.037% | 0.073% |
| **Lcas4** | 99.349% | 0.537% | 0.041% | 0.073% |
| **Lcas5** | 99.185% | 0.710% | 0.034% | 0.071% |
| **Overall** | **99.160%** | **0.724%** | **0.037%** | **0.079%** |

**Table S5.** Proportions of the *L. corsicanus* genome attributed to 4 different ancestries estimated with ELAI, in each analysed individual and overall.

| ind | *L. castroviejoi* | *L. granatensis* | *L. europaeus* | *L. timidus* |
|---|---|---|---|---|
| **Lcor1** | 99.888% | 0.013% | 0.016% | 0.039% |
| **Lcor2** | 99.903% | 0.015% | 0.015% | 0.033% |
| **Lcor3** | 99.894% | 0.019% | 0.017% | 0.034% |
| **Lcor4** | 99.859% | 0.016% | 0.056% | 0.033% |
| **Lcor5** | 99.896% | 0.020% | 0.017% | 0.030% |
| **Overall** | **99.888%** | **0.017%** | **0.024%** | **0.034%** |

**Table S6.** Average length of introgressed tracts (L) and the fraction of introgression (f) in *L. castroviejoi* (cas) from different origins (tim2cas – *L. timidus*; gra2cas – *L. granatensis*; eur2cas – *L. europaeus*), and in *L. granatensis* from *L. timidus* (tim2gra; recalculated from Seixas et al. (2018)). These estimates are presented for the complete datasets (Global) and for subsets representing Chromosomes Extremes or Centers. For the global analyses, statistics are shown merging tracts of *L. granatensis* and *L. timidus* origin in *L. castroviejoi* (gratim2cas) and estimates of introgression times are shown, considering the indicated recombination rate.

| **Event** | **Nr. Tracts** | **L (bp)** | **f** | **r (M/bp)** | **t (y)** |
|---|---|---|---|---|---|
| *Global* | | | | | |
| tim2cas | 92 | 25899 | 0.0006 | 1E-08 | 7729 |
| gra2cas | 484 | 47421 | 0.0054 | 1E-08 | 4454 |
| eur2cas | 20 | 36424 | 0.0002 | 1E-08 | 5494 |
| tim2gra | 2678 | 32939 | 0.0207 | 1E-08 | 6202 |
| gratim2cas | 90 | 46174 | 0.0060 | 1E-08 | 4359 |
| *Chromosome Extremes* | | | | | |
| tim2cas | 49 | 27860 | 0.0006 | - | - |
| gra2cas | 331 | 40728 | 0.0064 | - | - |
| eur2cas | 14 | 43762 | 0.0003 | - | - |
| tim2gra | 1614 | 31631 | 0.0239 | - | - |
| *Chromosome Centers* | | | | | |
| tim2cas | 43 | 23889 | 0.0005 | - | - |
| gra2cas | 151 | 61050 | 0.0043 | - | - |
| eur2cas | 6 | 19548 | 0.0001 | - | - |
| tim2gra | 1062 | 34556 | 0.0172 | - | - |

**Table S7.** Relative estimates of introgression times between events, estimated using introgression tracts inferred using ELAI, globally, and considering only chromosome extremes or centers. cas – *L. castroviejoi*; gra – *L. granatensis*; tim – *L. timidus*; eur – *L. castroviejoi*. For example, tim2cas indicates the timing of the event of admixture and introgression from *L. timidus* to *L. castroviejoi*.

| **Event 1** | **Event 2** | **Event 1 / Event 2** |
|---|---|---|
| *Global* | | |
| tim2cas | gra2cas | 1.74 |
| eur2cas | gra2cas | 1.23 |
| tim2cas | tim2gra | 1.25 |
| tim2gra | gra2cas | 1.39 |
| tim2gra | gratim2cas | 1.42 |
| *Chromosome Extremes* | | |
| tim2cas | gra2cas | 1.45 |
| eur2cas | gra2cas | 0.93 |
| tim2cas | tim2gra | 1.11 |
| tim2gra | gra2cas | 1.31 |
| *Chromosome Centers* | | |
| tim2cas | gra2cas | 2.54 |
| eur2cas | gra2cas | 3.11 |
| tim2cas | tim2gra | 1.42 |
| tim2gra | gra2cas | 1.79 |

**Table S8.** List of genes inspected in the Enrichment analysis for high introgression frequency of *L. granatensis* origin ($d_{XY}$ z-score outlier windows or showing *L. granatensis* introgression frequencies of at least 50% inferred in ELAI) and their function.

| Chr | Start | End | Gene name | Function |
|---|---|---|---|---|
| 1 | 15441317 | 15579207 | TSTD2 | |
| 1 | 15546196 | 15702310 | TDRD7 | biological regulation, regulation of biological process |
| 1 | 15634894 | 15687161 | ENSOCUG00000029006 | |
| 1 | 72045287 | 72127497 | NAA35 | biological regulation, regulation of biological process, regulation of cellular process |
| 1 | 72131198 | 72165745 | GOLM1 | biological regulation, regulation of biological process |
| 1 | 78359996 | 78439013 | SEMA4D | biological regulation, regulation of biological process, regulation of cellular process, cell communication, signaling, signal transduction |
| 1 | 79041952 | 79049783 | ENSOCUG00000022990 | biological regulation |
| 1 | 79074761 | 79146826 | SPIN1 | biological regulation, regulation of biological process, regulation of cellular process, cell communication, signaling, signal transduction |
| 1 | 98635213 | 98636307 | ENSOCUG00000038181 | |
| 1 | 100570515 | 100602524 | ENSOCUG00000038611 | |
| 1 | 147119401 | 147120476 | ENSOCUG00000028163 | |

| | | | | |
|---|---|---|---|---|
| 1 | 187613766 | 187614669 | ENSOCUG00000035290 | |
| 1 | 187693360 | 187694262 | ENSOCUG00000030075 | biological regulation, regulation of biological process, regulation of cellular process, cell communication, signaling, signal transduction |
| 1 | 187725519 | 187726448 | ENSOCUG00000024711 | biological regulation, regulation of biological process, regulation of cellular process, cell communication, signaling, signal transduction |
| 1 | 187738146 | 187739075 | ENSOCUG00000026679 | biological regulation, regulation of biological process, regulation of cellular process, cell communication, signaling, signal transduction |
| 1 | 187759150 | 187760076 | ENSOCUG00000008122 | biological regulation, regulation of biological process, regulation of cellular process, cell communication, signaling, signal transduction |
| 1 | 187790113 | 187791042 | ENSOCUG00000005268 | biological regulation, regulation of biological process, regulation of cellular process, cell communication, signaling, signal transduction |

| | | | | |
|---|---|---|---|---|
| 1 | 187803862 | 187804764 | ENSOCUG00000038037 | biological regulation, regulation of biological process, regulation of cellular process, cell communication, signaling, signal transduction |
| 1 | 187812242 | 187813171 | ENSOCUG00000039133 | biological regulation, regulation of biological process, regulation of cellular process, cell communication, signaling, signal transduction |
| 1 | 187828812 | 187829741 | ENSOCUG00000032013 | biological regulation, regulation of biological process, regulation of cellular process, cell communication, signaling, signal transduction |
| 1 | 187842009 | 187842938 | ENSOCUG00000021660 | biological regulation, regulation of biological process, regulation of cellular process, cell communication, signaling, signal transduction |
| 1 | 187854120 | 187855049 | ENSOCUG00000038724 | biological regulation, regulation of biological process, regulation of cellular process, cell communication, signaling, signal transduction |
| 1 | 187904739 | 187905656 | ENSOCUG00000033859 | biological regulation, regulation of biological process, regulation of |

| | | | | |
|---|---|---|---|---|
| | | | | cellular process, cell communication, signaling, signal transduction |
| 1 | 187925966 | 187927503 | ENSOCUG00000024521 | biological regulation, regulation of biological process, regulation of cellular process, cell communication, signaling, signal transduction |
| 1 | 189063513 | 189064460 | ENSOCUG00000036431 | biological regulation, regulation of biological process, regulation of cellular process, cell communication, signaling, signal transduction |
| 1 | 194587094 | 194599844 | ENSOCUG00000038329 | |
| 1 | 194597290 | 194599844 | ENSOCUG00000039622 | |
| 1 | 194599865 | 194731670 | KDM2A | biological regulation, regulation of biological process, regulation of cellular process |
| 1 | 194743802 | 194746827 | ENSOCUG00000038516 | |
| 1 | 194754486 | 194756335 | ENSOCUG00000029982 | |
| 1 | 194758430 | 194759825 | ENSOCUG00000006388 | |
| 1 | 194761747 | 194767569 | SSH3 | biological regulation, regulation of biological process, regulation of cellular process |
| 1 | 194770434 | 194835787 | RAD9A | biological regulation, regulation of biological process, regulation of cellular process, cell |

| | | | | |
|---|---|---|---|---|
| | | | | communication, signaling, signal transduction |
| 1 | 194802505 | 194804152 | ENSOCUG00000004005 | |
| 1 | 194811735 | 194819288 | CLCF1 | biological regulation, regulation of biological process, regulation of cellular process, cell communication, signaling, signal transduction |
| 2 | 308930 | 317348 | LYAR | biological regulation, regulation of biological process, regulation of cellular process |
| 2 | 94508442 | 94611216 | ENSOCUG00000028202 | |
| 2 | 98129481 | 98247972 | ENSOCUG00000029292 | |
| 2 | 98223558 | 98454000 | ENSOCUG00000027394 | |
| 2 | 98926095 | 98926409 | ENSOCUG00000025374 | |
| 2 | 98961851 | 98993640 | ENSOCUG00000006530 | |
| 2 | 100020636 | 100144983 | REEP1 | |
| 2 | 100147102 | 100162537 | MRPL35 | |
| 2 | 100167214 | 100212489 | IMMT | biological regulation |
| 2 | 100217116 | 100247027 | ENSOCUG00000010161 | |
| 2 | 100218424 | 100218558 | SNORD94 | |
| 2 | 100247451 | 100326763 | POLR1A | biological regulation, regulation of biological process, regulation of cellular process |
| 2 | 100440078 | 100505486 | ST3GAL5 | |
| 2 | 100543944 | 100553597 | ATOH8 | biological regulation, regulation of biological process, regulation of |

| | | | | |
|---|---|---|---|---|
| | | | | cellular process, cell communication, signaling, signal transduction |
| 3 | 32469752 | 32481040 | ENSOCUG00000035396 | |
| 3 | 32483817 | 32584463 | ENSOCUG00000024432 | |
| 3 | 32505854 | 32510754 | ENSOCUG00000028024 | |
| 3 | 32574069 | 32585184 | ZNF300 | biological regulation, regulation of biological process, regulation of cellular process |
| 3 | 54822318 | 54831487 | ENSOCUG00000031970 | |
| 3 | 54842765 | 54910760 | CPEB4 | biological regulation, regulation of biological process, regulation of cellular process |
| 3 | 146490086 | 146740453 | ADCY8 | biological regulation, regulation of biological process, regulation of cellular process, cell communication, signaling, signal transduction |
| 3 | 148923207 | 148936788 | CCN4 | biological regulation, regulation of biological process, regulation of cellular process, cell communication, signaling, signal transduction |
| 4 | 9109254 | 9148757 | ENSOCUG00000023941 | |
| 4 | 19488979 | 19532701 | ENSOCUG00000030426 | |

| | | | | |
|---|---|---|---|---|
| 4 | 22560026 | 22963461 | TASP1 | biological regulation, regulation of biological process, regulation of cellular process |
| 4 | 24276445 | 24309484 | BTBD3 | |
| 4 | 28349814 | 28509022 | HAO1 | |
| 4 | 37189666 | 37226242 | SP1 | biological regulation, regulation of biological process, regulation of cellular process |
| 4 | 37775877 | 37775984 | MIR196A2 | |
| 4 | 37784456 | 37787223 | HOXC9 | biological regulation, regulation of biological process, regulation of cellular process |
| 4 | 37793427 | 37795810 | HOXC8 | biological regulation, regulation of biological process, regulation of cellular process |
| 4 | 38080742 | 38101057 | COPZ1 | |
| 4 | 38111631 | 38112818 | ENSOCUG00000010753 | biological regulation, regulation of biological process, regulation of cellular process, cell communication, signaling, signal transduction |
| 4 | 38121032 | 38141067 | ZNF385A | biological regulation, regulation of biological process, regulation of cellular process, cell communication, signaling, signal transduction |

| | | | | |
|---|---|---|---|---|
| 4 | 38207618 | 38289482 | NCKAP1L | biological regulation, regulation of biological process, regulation of cellular process, cell communication, signaling, signal transduction |
| 4 | 38294892 | 38323526 | PDE1B | biological regulation, regulation of biological process, regulation of cellular process, cell communication, signaling, signal transduction |
| 4 | 38323744 | 38332651 | ENSOCUG00000013555 | biological regulation, regulation of biological process, regulation of cellular process, cell communication, signaling, signal transduction |
| 4 | 43108831 | 43197947 | C12orf56 | |
| 4 | 48184819 | 48214405 | CPSF6 | biological regulation, regulation of biological process, regulation of cellular process |
| 4 | 48613716 | 48618906 | FRS2 | biological regulation, regulation of biological process, regulation of cellular process, cell communication, signaling, signal transduction |
| 4 | 49389824 | 49465807 | KCNMB4 | biological regulation, cell communication, signaling, |
| 4 | 51397572 | 51830973 | TRHDE | |

| | | | | |
|---|---|---|---|---|
| 4 | 60062265 | 60210781 | LIN7A | biological regulation, cell communication, signaling, |
| 4 | 60587600 | 61132121 | PPFIA2 | biological regulation, regulation of biological process, regulation of cellular process |
| 4 | 61629400 | 61741612 | METTL25 | |
| 4 | 62760057 | 62775626 | ENSOCUG00000035502 | |
| 4 | 67196551 | 67242936 | TMTC3 | biological regulation, regulation of biological process, regulation of cellular process |
| 4 | 67493134 | 67575339 | KITLG | biological regulation, regulation of biological process, regulation of cellular process, cell communication, signaling, signal transduction |
| 4 | 75474542 | 75511662 | CDK17 | |
| 4 | 78037409 | 79260279 | ANKS1B | |
| 5 | 13310566 | 13386213 | PSME3IP1 | biological regulation, regulation of biological process, regulation of cellular process |
| 5 | 13386255 | 13439299 | RSPRY1 | |
| 5 | 13449903 | 13471380 | PLLP | |
| 5 | 22005555 | 22036447 | ENSOCUG00000026523 | |
| 5 | 22041880 | 22048898 | CMTM3 | biological regulation, regulation of biological process, regulation of cellular process, cell |

| | | | | |
|---|---|---|---|---|
| | | | | communication, signaling, signal transduction |
| 5 | 22055909 | 22167329 | DYNC1LI2 | |
| 5 | 22171617 | 22212781 | TERB1 | |
| 5 | 22204313 | 22204506 | U3 | |
| 5 | 22222310 | 22247009 | NAE1 | biological regulation, regulation of biological process, regulation of cellular process, cell communication, signaling, signal transduction |
| 6 | 1401914 | 1769725 | RBFOX1 | biological regulation, regulation of biological process, regulation of cellular process |
| 6 | 3975325 | 4009191 | ENSOCUG00000034442 | |
| 6 | 5390743 | 5869252 | SNX29 | |
| 6 | 8547405 | 8640042 | VPS35L | |
| 6 | 8644270 | 8663005 | CCP110 | biological regulation, regulation of biological process, regulation of cellular process |
| 6 | 13887348 | 13974482 | USP31 | |
| 7 | 1382865 | 1528527 | PTN | biological regulation, regulation of biological process, regulation of cellular process, cell communication, signaling, signal transduction |
| 7 | 1578626 | 2071619 | DGKI | biological regulation, regulation of biological process, regulation of |

| | | | | |
|---|---|---|---|---|
| | | | | cellular process, cell communication, signaling, signal transduction |
| 7 | 9252482 | 9334638 | COPG2 | |
| 7 | 11213822 | 11547011 | ENSOCUG00000014032 | |
| 7 | 11630125 | 12818701 | LRGUK | |
| 7 | 14474543 | 14551958 | KLHDC10 | |
| 7 | 14551979 | 14600560 | ZC3HC1 | biological regulation, regulation of biological process, regulation of cellular process, cell communication, signaling, signal transduction |
| 7 | 15080461 | 15128956 | STRIP2 | biological regulation, regulation of biological process |
| 7 | 67114365 | 68644978 | DPP10 | biological regulation, regulation of biological process, regulation of cellular process |
| 7 | 103860153 | 103987125 | GRB14 | biological regulation, regulation of biological process, regulation of cellular process, cell communication, signaling, signal transduction |
| 7 | 104038187 | 104251392 | COBLL1 | |
| 7 | 104321307 | 104385124 | SLC38A11 | |
| 7 | 104351687 | 104351816 | ENSOCUG00000019821 | |
| 7 | 104491906 | 104607488 | SCN3A | voltage-gated sodium channel complex |

| | | | | |
|---|---|---|---|---|
| 7 | 104665799 | 104796938 | SCN2A | biological regulation, regulation of biological process, regulation of cellular process, cell communication, signaling, signal transduction, voltage-gated sodium channel complex |
| 7 | 104824257 | 104850959 | ENSOCUG00000032787 | |
| 7 | 104892048 | 105087752 | CSRNP3 | biological regulation, regulation of biological process, regulation of cellular process |
| 7 | 105115129 | 105115596 | ENSOCUG00000022341 | |
| 7 | 105155110 | 105182854 | GALNT3 | |
| 7 | 105247455 | 105396969 | TTC21B | biological regulation, regulation of biological process, regulation of cellular process, cell communication, signaling, signal transduction |
| 7 | 105415056 | 105494742 | SCN1A | biological regulation, cell communication, signaling, voltage-gated sodium channel complex |
| 7 | 105535714 | 105535820 | U6 | |
| 8 | 6368815 | 6547738 | KIF21A | |
| 8 | 8903444 | 9118144 | BICD1 | biological regulation, regulation of biological process, regulation of cellular process |
| 8 | 9219782 | 9249780 | RESF1 | |

| | | | | |
|---|---|---|---|---|
| 8 | 9272067 | 9272606 | ENSOCUG00000033986 | |
| 8 | 9423421 | 9470014 | AMN1 | |
| 8 | 9454712 | 9454828 | U5 | |
| 8 | 9480156 | 9484835 | ENSOCUG00000023987 | |
| 8 | 9538809 | 9741935 | DENND5B | biological regulation, regulation of biological process |
| 8 | 37314705 | 37492177 | KDM5A | biological regulation, regulation of biological process, regulation of cellular process |
| 8 | 37503424 | 37503678 | ENSOCUG00000023846 | |
| 8 | 37509673 | 37613485 | IQSEC3 | biological regulation, regulation of biological process, regulation of cellular process, cell communication, signaling, signal transduction |
| 9 | 1885725 | 2145078 | ENSOCUG00000035357 | |
| 9 | 3603317 | 3925918 | MYRIP | |
| 9 | 3899728 | 3901720 | ENSOCUG00000029704 | |
| 9 | 6755107 | 6847682 | ENSOCUG00000036497 | |
| 9 | 6938543 | 7007704 | KBTBD12 | |
| 9 | 7034415 | 7072608 | SEC61A1 | |
| 9 | 7044984 | 7185439 | RUVBL1 | biological regulation, regulation of biological process, regulation of cellular process, cell communication, signaling, signal transduction |
| 9 | 87502229 | 87513280 | ATP5F1A | |

| | | | | |
|---|---|---|---|---|
| 9 | 87517360 | 87536403 | HAUS1 | |
| 9 | 93286394 | 94114886 | DCC | biological regulation, regulation of biological process, regulation of cellular process, cell communication, signaling |
| 10 | 15934034 | 16473294 | PDE1C | biological regulation, regulation of biological process, regulation of cellular process, cell communication, signaling, signal transduction |
| 10 | 19832513 | 19882325 | KIAA0895 | |
| 10 | 21116879 | 21121656 | GPR141 | biological regulation, regulation of biological process, regulation of cellular process, cell communication, signaling, signal transduction |
| 10 | 21147133 | 21148022 | ENSOCUG00000024486 | |
| 10 | 21176130 | 21219208 | ENSOCUG00000015861 | |
| 10 | 21235086 | 21243552 | SFRP4 | biological regulation, regulation of biological process, regulation of cellular process, cell communication, signaling, signal transduction |
| 10 | 21247907 | 21270997 | EPDR1 | |
| 10 | 21480758 | 21481093 | ENSOCUG00000036137 | |
| 11 | 40285367 | 40739663 | FBXL7 | |
| 11 | 56206706 | 56208844 | ENSOCUG00000029955 | |

| | | | | |
|---|---|---|---|---|
| 11 | 56221803 | 56227231 | RAD1 | biological regulation, regulation of biological process, regulation of cellular process, cell communication, signaling, signal transduction |
| 11 | 56320259 | 56362897 | AGXT2 | biological regulation, regulation of biological process, regulation of cellular process |
| 11 | 56363806 | 56565013 | PRLR | biological regulation, regulation of biological process, regulation of cellular process, cell communication, signaling, signal transduction |
| 12 | 9745147 | 9746083 | ENSOCUG00000022359 | |
| 12 | 23019284 | 23048090 | ENSOCUG00000006769 | |
| 12 | 23039791 | 23041269 | ENSOCUG00000025822 | |
| 12 | 39165077 | 39169010 | ENSOCUG00000035733 | |
| 12 | 39171952 | 39172977 | ENSOCUG00000032803 | |
| 12 | 39173820 | 39175967 | ENSOCUG00000039348 | |
| 12 | 72763182 | 72921498 | ENSOCUG00000032043 | |
| 12 | 75957741 | 75966862 | SRSF12 | biological regulation, regulation of biological process, regulation of cellular process |
| 12 | 126999350 | 127245281 | MAP3K5 | biological regulation, regulation of biological process, regulation of cellular process, cell |

| | | | | |
|---|---|---|---|---|
| | | | | communication, signaling, signal transduction |
| 12 | 127368196 | 127406612 | SLC35D3 | biological regulation, regulation of biological process, regulation of cellular process |
| 13 | 7001249 | 7137678 | CEP350 | |
| 13 | 15260025 | 16352709 | DPP6 | biological regulation, regulation of biological process, regulation of cellular process |
| 13 | 31095033 | 31311165 | ATF6 | biological regulation, regulation of biological process, regulation of cellular process, cell communication, signaling, signal transduction |
| 13 | 32394249 | 32507941 | ENSOCUG00000022552 | |
| 13 | 32433771 | 32470445 | SLAMF1 | biological regulation, regulation of biological process, regulation of cellular process, cell communication, signaling, signal transduction |
| 13 | 34654583 | 34655527 | ENSOCUG00000001639 | biological regulation, regulation of biological process, regulation of cellular process, cell communication, signaling, signal transduction |
| 13 | 34670818 | 34678508 | ENSOCUG00000022281 | biological regulation, regulation of biological process, regulation of |

| | | | | |
|---|---|---|---|---|
| | | | | cellular process, cell communication, signaling, signal transduction |
| 13 | 34954808 | 35073530 | ENSOCUG00000008307 | |
| 13 | 37158060 | 37335037 | ENSOCUG00000034353 | |
| 13 | 37159192 | 37374813 | ASH1L | biological regulation, regulation of biological process, regulation of cellular process, cell communication, signaling, signal transduction |
| 13 | 41805373 | 41821930 | ENSOCUG00000022805 | |
| 13 | 47077538 | 47138353 | VTCN1 | biological regulation, regulation of biological process, regulation of cellular process |
| 13 | 47169788 | 47178215 | TRIM45 | |
| 13 | 51967998 | 51969299 | ENSOCUG00000023372 | |
| 13 | 66456505 | 67430660 | DPYD | |
| 13 | 75695873 | 75851300 | KYAT3 | |
| 13 | 75844666 | 75999099 | PKN2 | biological regulation, regulation of biological process, regulation of cellular process, cell communication, signaling, signal transduction |
| 13 | 77712346 | 77910608 | ENSOCUG00000010672 | |
| 13 | 80727235 | 80949545 | PRKACB | biological regulation, regulation of biological process, regulation of cellular process, cell |

| | | | | |
|---|---|---|---|---|
| | | | | communication, signaling, signal transduction |
| 13 | 87597110 | 87703434 | MIGA1 | |
| 13 | 87714216 | 87784995 | USP33 | biological regulation, regulation of biological process, regulation of cellular process, cell communication, signaling, signal transduction |
| 13 | 99337246 | 99424659 | DNAI4 | |
| 13 | 101596434 | 101674992 | RAVER2 | |
| 13 | 103843429 | 104078176 | DOCK7 | biological regulation, regulation of biological process, regulation of cellular process, cell communication, signaling, signal transduction |
| 13 | 113052119 | 113282809 | GLIS1 | biological regulation, regulation of biological process, regulation of cellular process |
| 13 | 113322494 | 113329720 | ENSOCUG00000001341 | biological regulation, regulation of biological process, regulation of cellular process |
| 13 | 113374888 | 113375898 | ENSOCUG00000030578 | |
| 13 | 113475513 | 113539335 | LRP8 | biological regulation, regulation of biological process, regulation of cellular process, cell communication, signaling, signal transduction |

| | | | | |
|---|---|---|---|---|
| 13 | 113549246 | 113556839 | MAGOH | biological regulation, regulation of biological process, regulation of cellular process |
| 13 | 113553616 | 113553722 | U6 | |
| 13 | 113560288 | 113566628 | CZIB | |
| 13 | 113576071 | 113595304 | CPT2 | biological regulation, regulation of biological process |
| 13 | 113579850 | 113579949 | U6 | |
| 13 | 114146858 | 114287036 | TUT4 | |
| 13 | 124058910 | 124223727 | CCDC30 | |
| 13 | 124222520 | 124244815 | ZMYND12 | |
| 13 | 124261082 | 124290797 | RIMKLA | |
| 13 | 129356566 | 129370185 | SRSF10 | biological regulation, regulation of biological process, regulation of cellular process |
| 13 | 129373633 | 129376775 | ENSOCUG00000038691 | biological regulation, regulation of biological process |
| 13 | 131935886 | 132330532 | EIF4G3 | |
| 13 | 132321361 | 132321521 | U1 | |
| 13 | 135109985 | 135262202 | PUM1 | biological regulation, regulation of biological process, regulation of cellular process, cell communication, signaling, signal transduction |
| 13 | 135244323 | 135244407 | ENSOCUG00000021668 | |

| | | | | |
|---|---|---|---|---|
| 13 | 136462610 | 136587222 | EPB41 | biological regulation, regulation of biological process, regulation of cellular process |
| 13 | 137721511 | 137752045 | FAM76A | |
| 14 | 691101 | 762004 | PIK3R4 | biological regulation, regulation of biological process, regulation of cellular process, cell communication |
| 14 | 737715 | 737842 | 5S_rRNA | |
| 14 | 761497 | 934532 | COL6A6 | |
| 14 | 2341676 | 2648399 | RFTN1 | biological regulation, regulation of biological process, regulation of cellular process, cell communication, signaling, signal transduction |
| 15 | 4030371 | 4238133 | CSGALNACT1 | |
| 15 | 12609533 | 12611770 | ENSOCUG00000035083 | |
| 15 | 12618216 | 12622501 | ENSOCUG00000031102 | |
| 15 | 18173522 | 18459851 | SLC10A7 | biological regulation |
| 15 | 20734409 | 20789695 | FREM3 | |
| 15 | 20841758 | 20883541 | SMARCA5 | biological regulation, regulation of biological process, regulation of cellular process, , , |
| 15 | 20961381 | 21092482 | GAB1 | biological regulation, regulation of biological process, regulation of cellular process, cell |

| | | | | |
|---|---|---|---|---|
| | | | | communication, signaling, signal transduction |
| 15 | 43060237 | 43171532 | ENSOCUG00000038394 | |
| 17 | 80340533 | 80519441 | ATP10A | biological regulation, regulation of biological process, regulation of cellular process |
| 18 | 3020835 | 3026299 | ZNF25 | biological regulation, regulation of biological process, regulation of cellular process |
| 18 | 3084222 | 3101792 | ENSOCUG00000029543 | biological regulation, regulation of biological process, regulation of cellular process |
| 18 | 3172108 | 3247233 | ENSOCUG00000029443 | biological regulation, regulation of biological process, regulation of cellular process |
| 18 | 6522135 | 6591232 | ZFAND4 | |
| 18 | 7460895 | 8191494 | GRID1 | |
| 18 | 42383100 | 42989156 | ENSOCUG00000015280 | |
| 18 | 61065683 | 61260227 | NRAP | |
| 19 | 4124261 | 4158706 | LRRC75A | |
| 19 | 4160060 | 4190795 | ENSOCUG00000033937 | |
| 19 | 31133263 | 31142229 | ENSOCUG00000011338 | |
| 19 | 43820936 | 43825782 | TMEM106A | biological regulation, regulation of biological process, regulation of cellular process, cell communication, signaling, signal transduction |

| | | | | |
|---|---|---|---|---|
| 19 | 43829927 | 43835288 | ENSOCUG00000033652 | |
| 19 | 43837731 | 43837921 | U2 | |
| 19 | 43838409 | 43864540 | ENSOCUG00000030666 | |
| 19 | 43838431 | 43838537 | U6 | |
| 19 | 43849321 | 43849498 | U2 | |
| 19 | 43990472 | 44044059 | DHX8 | |
| 19 | 51999202 | 52088717 | RGS9 | biological regulation, regulation of biological process, regulation of cellular process, cell communication, signaling, signal transduction |
| 19 | 52117642 | 52119972 | ENSOCUG00000036189 | |
| 19 | 52119221 | 52160069 | GNA13 | biological regulation, regulation of biological process, regulation of cellular process, cell communication, signaling, signal transduction |
| 19 | 56088008 | 56149875 | ENSOCUG00000031100 | |
| 20 | 8972192 | 9159086 | ENSOCUG00000015859 | biological regulation, regulation of biological process, regulation of cellular process, cell communication, signaling, signal transduction |
| 20 | 10905320 | 10996757 | RHOJ | biological regulation, regulation of biological process, regulation of cellular process, cell |

| | |
|---|---|
| | communication, signaling, signal transduction |
